# Effect on the conformations of spike protein of SARS-CoV-2 due to mutation

**DOI:** 10.1101/2022.05.11.491583

**Authors:** Aayatti Mallick Gupta, Jaydeb Chakrabarti

## Abstract

The spike protein of SARS CoV-2 mediates receptor binding and cell entry and is the key immunogenic target for virus neutralization and the present attention of many vaccine layouts. It exhibits significant conformational flexibility. We study the structural fluctuations of spike protein among the most common mutations appeared in variant of concerns (VOC). We report the thermodynamics of conformational changes in mutant spike protein with respect to the wildtype from the distributions of the dihedral angles obtained from the equilibrium configurations generated via all-atom molecular dynamics simulations. We find that the mutation causes the increase in distance between N-terminal domain and receptor binding domain leading to an obtuse angle cosine *θ* distribution in the trimeric structure in spike protein. Thus, increase in open-state is conferred to the more infectious variants of SARS-CoV-2. The thermodynamically destabilized and disordered residues of receptor binding motif among the mutant variants of spike protein are proposed to serve as better binding sites for host factor. We identify a short stretch of region connecting the N-terminal domain and receptor binding domain forming linker loop where many residues undergo stabilization in the open state compared to the closed one.

## 1. Introduction

Since the first documented cases of SARS-CoV-2^1^ infection in Wuhan, China in late 2019^2^, the COVID-19 pandemic poses an unprecedented threat to the global public health, with more than 286 million infections and over 5.4 million deaths around the world (https://www.who.int/emergencies/diseases/novel-coronavirus-2019/situation-reports/). Despite rapid development and emergency authorization of vaccines, immune escape mutants have emerged, and SARS-CoV-2 infections remain a concern for the global community. Although vaccination has significantly lowered the rates of hospitalization, severity and death^3–6^, current vaccines do not confer absolute prevention of upper airway transmission of SARS-CoV-2. The numbers of vaccine breakthrough infections and re-infections, consequently, have been continuously reported^7–9^. It is of vital importance to examine the impact of the mutant variants as soon as they are detected by genomic sequence analysis. In the light of crucial role of spike protein in virus infection and host immune evasion, studies have been prioritized on the emerging mutations of spike protein circulating SARS-CoV-2 strains and investigations on their biological significance^10^. The structural information is essential for structure-based design of vaccine immunogens and entry inhibitors of SARS-CoV-2.

Spike protein (180–200 kDa) of SARS-CoV-2 virus consists of a homo-trimeric large clover-shaped protrusion that mediates viral entry to the host cell through the human ACE2 receptor^11^ distributed mainly in the lung, intestine, heart, and kidney, and alveolar epithelial type II cells^12^. Each spike monomer (1273 aa) consists of a signal peptide located at the N-terminus, the S1 subunit and the S2 subunit. The S1 subunit is responsible for receptor binding comprise of an N-terminal domain (NTD) and a receptor-binding domain (RBD). A short stretch of amino acid residues connects the NTD arm with that of the RBD forms the linker. The S2 subunit comprises of the fusion peptide (FP), heptapeptide repeat sequence 1 (HR1), heptapeptide repeat sequence 2 (HR2), transmembrane (TM) domain and cytoplasm domain. The S2 domain entangles to create the stalk, transmembrane, and small intracellular domains^13^. Spike protein subsists in a metastable, prefusion conformation acting as an inactive precursor. However, when the virus interacts with the host cell, extensive structural rearrangement of the spike protein occurs, by cleaving it into S1 and S2 subunits, allowing the virus to fuse with the host cell membrane. RBD located in the S1 subunit interacts with the cell receptor ACE2 in the region of aminopeptidase N. Remarkable conformational heterogeneity can be found in the RBD region. Within a single protomer, the RBD could adopt a receptor inaccessible closed ‘down’ state in which the RBD is buried at the interface between the protomers and be accessible to ACE2, or an open ‘up’ state that enables exposure of the receptor-binding motif which mediates interaction with ACE2^14,15^.

During late 2020, the emergence of variants that posed an increased risk to global public health prompted WHO for the characterization of specific Variants of Interest (VOIs) and Variants of Concern (VOCs) of SARS-CoV-2. Currently, there are five VOCs: Alpha variant (B.1.1.7; RBD mutations: N501Y, A570D), Beta variant (B.1.351; RBD mutations: K417N, E484K, and N501Y), Gamma variant (P.1, B.1.1.28.1; RBD mutations: K417N/T, E484K, and N501Y), Delta variant (B.1.617.2; RBD mutations: L452R, T478K) and Omicron variant (B. 1.1.529; multiple RBD mutations; new VOC designated on 26^th^ Nov, 2021). The mutated variants are characterized by the presence of genetic changes which are known to affect virus characteristics such as transmissibility, disease severity, immune escape, and diagnostic or therapeutic escape^16–23^. It has been observed that alpha, beta and gamma variants contain some common mutations, like E484K and N501Y in the receptor binding motif of RBD region. N501Y and E484K have been reported for increase in ACE2 binding affinity^24^ and decrease in efficacy for antibody binding accountable for immune evasion^25,26^. Thus, particularly the RBD variants in SARS-CoV-2 are vital to recognize mutant viral strains with higher transmissibility and the potentiality to bring about immune invasion. Here we study the effect of mutations in RBD on the conformations of spike protein of SARS-CoV-2.

We are particularly interested in the stability of the mutated protein with respect to the wild type. The relative stability of protein conformations has been extracted from mean field description based on conformational thermodynamics data^27–30^. In this method, the changes in thermodynamics free energy and entropy of a protein in a conformation with respect to a reference conformation are estimated from fluctuations of the dihedral angles in the two states over the simulated trajectories. Earlier studies based on conformational thermodynamics suggest that the destabilized and disordered residues of a protein in a particular conformation are the functional ones in that state, leading to binding specificity^30^.

We perform all atom MD simulations of the complete spike protein of SARS-CoV-2 in its wild type and the several mutated VOC strains, like K417N, L452R, E484K and N501Y using the GROMOS96 53a6 force-field in the GROMACS 2018.6 package. we find that the spike protein prefers to be in open state under mutations in the RBD, while a close conformation is preferred in the wild type. We calculate the thermodynamics cost of conformational changes. The RDB residues in mt(K417N)-spike, mt(L452R)-spike, mt(E484K)-spike, mt(N501Y)-spike and mt(mult)-spike show destabilization and disorder with respect to the wild type conformation. On the other hand, the NTD region does not reveal much significant changes in stability and order. The linker loop reveals increase in order and stability in the mutated variants. Our studies may shade important light to the virulence of the viral species.

## 2. Materials and Methods

### 2.1. System Preparation and Simulation details

The cryo EM structure of SARS CoV-2 spike trimer in a tightly closed state (7DF3) as well as in its open state (7DK3) are considered to study the conformational dynamics of spike protein and its effect on mutation. The mutations are obtained from the cryo EM structure. We have considered the most common mutations appeared in VOC strains like K417N, L452R, E484K and N501Y. We have also chosen a system which contains multiple mutations together designated as ‘mt(mult)-spike’ and a separate system which include only H69del and V70del without any other spike mutations as ‘mt(del)-spike’.

We perform 1 μs-long all-atom MD simulation using the standard protocol for isothermal isobaric ensemble (NPT) with 310 K and 1-atm pressure in GROMACS^31^ package. We use periodic boundary conditions, spc216 water model and GROMOS9353a6^32^ force field for simulations in GROMACS 2018.6 package. Electro-neutrality is maintained by adding monovalent ions Na+ and Cl−. Long ranged columbic interactions are considered using PME approach^33^. LINCS algorithm^34^ is used to constraint the bonds and leap-frog integration is used to perform simulation. Minimization is done for 50,000 steps using the steepest descent algorithms. Equations of motion are integrated using leap-frog algorithm with an integration time step of 2fs. Systems are equilibrated through 2 steps (NVT & NPT) using position restraints to heavy atoms. NVT and NPT equilibration is carried out at 300K Temperature and 1 Bar pressure. We maintain the total number of particles (N = 166251), pressure and temperature same for all the systems to make the simulated ensembles equivalent.

Seven independent MD simulations for wt-spike, mt(K417N)-spike, mt(L452R)-spike, mt(E484K)-spike, mt(N501Y)-spike, mt(mult)-spike and mt(del)-spike systems are performed for 1 μs with different initial conditions (i) open conformation and (ii) closed conformation. All runs are repeated three times with different seeds. The equilibrations of the simulated structures are assessed from the saturation of the root mean squared deviations, RMSD. All the data have been averaged over six independent trajectories for each system.

### 2.2 Conformational thermodynamics

Conformational thermodynamics changes for spike proteins and its mutated varieties at different conformations are estimated properly from equilibrium fluctuations of the dihedral angles using the Histogram Based Method (HBM)^30^. Equilibrium conformational changes in free energy is defined by 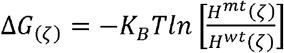 where *H^mt^*(*ζ*) and *H^wt^*(*ζ*)signify peak value of normalized probability distribution of protein dihedral (*ζ*) of mutated and wildtype spike protein respectively, and *K_B_*; the Boltzmann’s Constant. Conformational entropy change associated with a particular dihedral (*ζ*) at a temperature T is calculated using *T*Δ*S* = *T*(*S^mt^*(*ζ*) - *S^wt^*(*ζ*))where *S^mt^*(*ζ*) and *S^wt^*(*ζ*) can be obtained using Gibbs entropy formula *S*(*ζ*) = –*K_B_T* ∑_*i*_*H_i_*(*ζ*) *InH_i_*(*ζ*); sum is taken over histogram bins.

## 3. Results and discussion

### 3.1. Effect of mutation on spike protein conformation

In the current study we have focused only into the S1 subunit that interacts with the host cell receptor ACE2, consisting of the NTD (14-306 residues), the RBD (331-528 residues)and the linker (306-331). The crystal structure RBD of spike protein can be in both closed (fig. 1(a))and open (fig. 1(b)) conformations. We show equilibrium snapshots for different cases in the figure 1. We have found that the wt-spike protein attains a closed conformation upon all cases (fig. 1(c)). However, in all the mutated systems, mt(K417N)-spike, mt(L452R)-spike, mt(E484K)-spike, mt(N501Y)-spike and mt(mult)-spike, the open conformation is observed (fig1 (d)-(h)). On the other hand, we have observed that the mt(del)-spike protein prefers close conformation (fig. 1(i)).

**Fig1:**
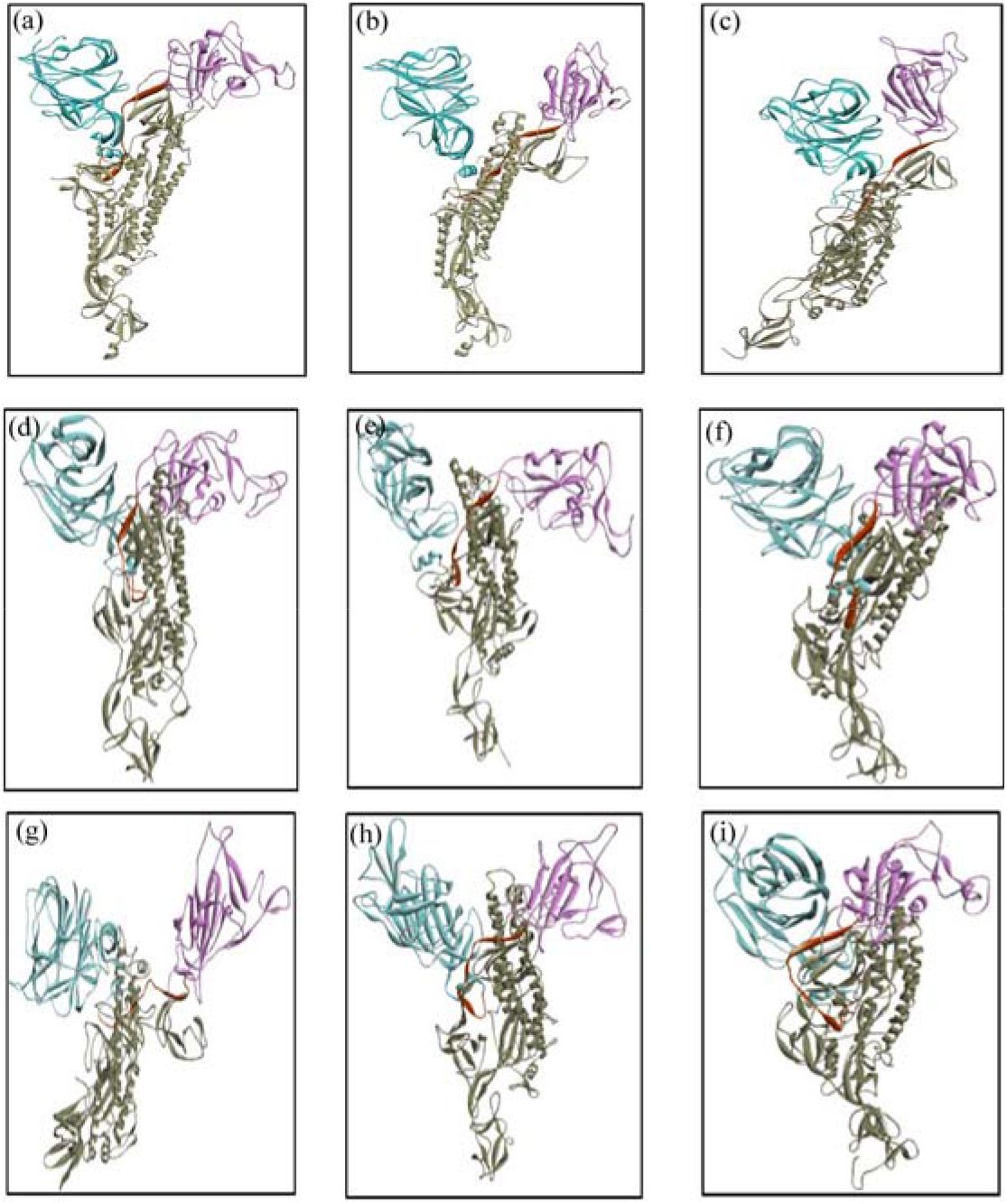
Color online: Structure of trimeric spike protein of SARS-CoV-2 in its wildtype and mutated form at different conformation states. NTD is shown in blue, RBD in pink and Linker arm in brown. (a) spike protein at its closed state (PDB id: 7DF3). (b) Open state of spike protein (PDB id: 7DY3). Snapshot at 1μs time span of (c) wt-spike (d) mt(K417N)-spike (e) mt(L452R)-spike (f) mt(E484K)-spike (g) mt(N501Y)-spike (h) mt(mult)-spike (i) mt(del)-spike.

To quantify the conformational changes, we consider the centers of mass for NTD and RBD region respectively from the equilibrated trajectory of each of the system and then the distance *S* between the centers of mass of NTD and the RBD arm has been calculated over simulated trajectories. We show the distribution of *S*, *H*(*S*) over the equilibrium trajectories for all the systems in Fig. 2a. For the wt-spike protein, *H*(*S*) shows a sharp peak at around 3 nm and mt(del)-spike shows approximately around 3.5 nm. This distance is comparable to the crystal structure data. However, the rest all of the mutant variants show peak in the range 4.5 nm to 6 nm. Thus, the distance between the NTD and RBD of the spike protein increases due to mutation, so that the RBD prefers the open conformation to the closed one. We further consider the cosine of the angle *θ* between the vectors joining center of mass of NTD arm, those of the linker and the RBD arm. We show the distribution of cos*θ*, *H*(*cosθ*) over the equilibrium trajectories for all the systems in Fig. 2b. Both wt-spike and mt(del)-spike show peak around *θ*=90°. In mt(K417N)-spike there is a peak at *θ*= 110°, mt(N501Y)-spike confers peak at *θ*= 100° and mt(E484K)-spike shows peak at *θ*= 105°. mt(L452R)-spike shows a shift in peak for wt-spike at *θ*= 180°, while in the mt(mult)-spike maximum peak can be found at *θ*= 120°. This data also supports open conformation in mutant variants of SARS-CoV-2.

**Fig2:**
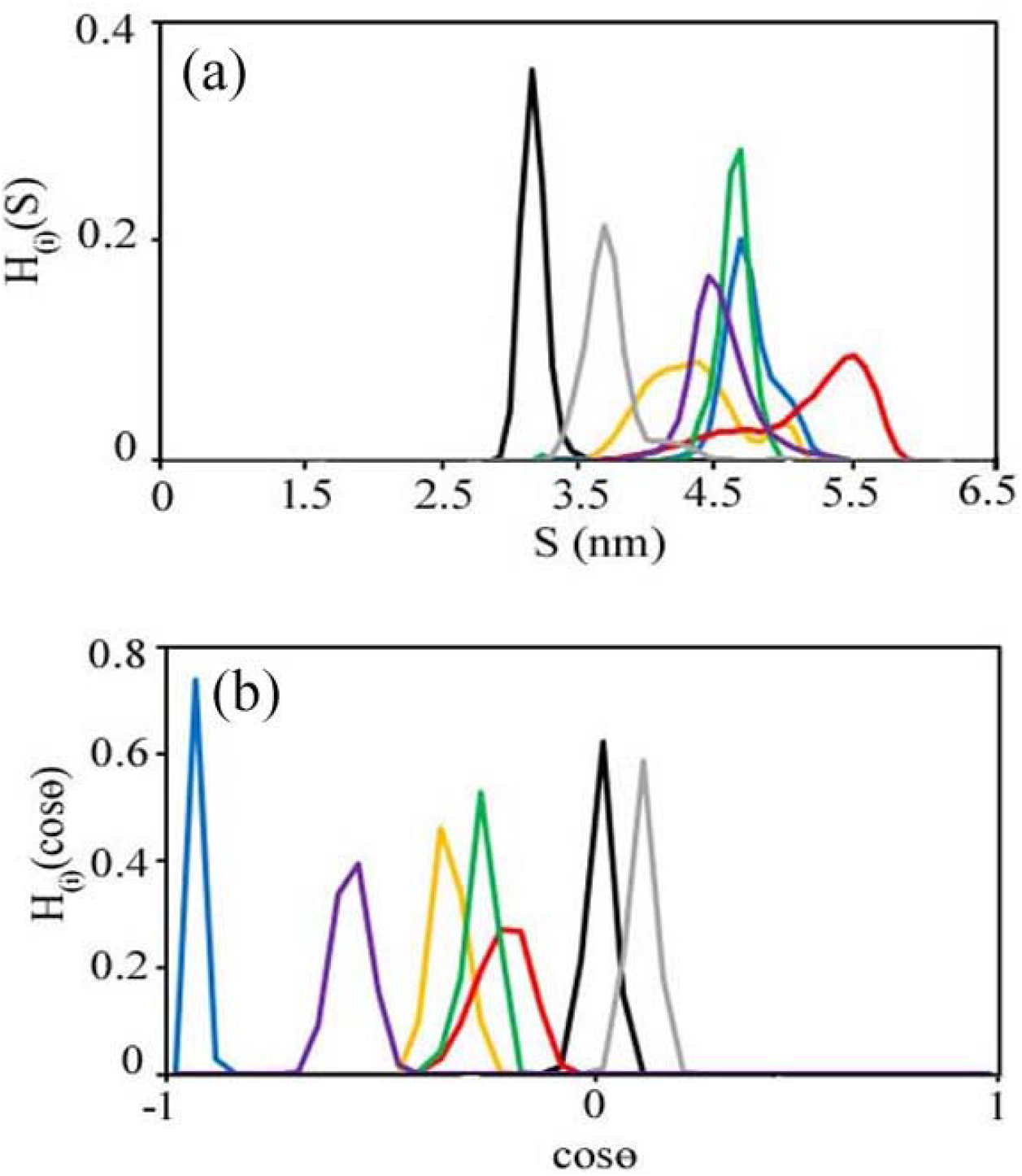
Color online: (a) Histogram distribution of the distance between the *C_α_* atoms between the two center of mass of NTD arm to that of RBD arm calculated over time span. S, H(S) over the equilibrium trajectories for all the systems are represented by wt-spike in black, mt(del)-spike in grey, mt(K417N)-spike in yellow, mt(L452R)-spike in blue, mt(E484K)-spike in green, mt(N501Y)-spike in red and mt(mult)-spike in violet. (b) The distribution of cos*θ*, H(cos*θ*) over the equilibrium trajectories for all the systems (color demarcation are same as (a)). Mutation causes the RBD arm to be more distantly apart from NTD arm, causing an obtuse angle cosine *θ* distribution in the trimeric structure in spike protein.

### 3.2 Conformational thermodynamics due to mutation

The flexibility of the protein conformations is given in terms of the dihedral fluctuations. The dihedral angles φ, ψ of the backbone and χ_1_ of side chain for different region of the spike proteins in wt-spike and variant systems are computed from the equilibrated portion of the trajectories. We denote the distribution of a dihedral angle *θ*(either of φ, ψ and χ_1_) of the *i*-th residue in wt-spike by 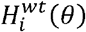,and variant systems by 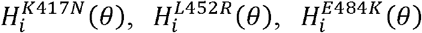, 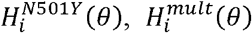 and 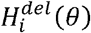, depending on the kind of modification over the wt-spike protein. A broadened or multiple-peaked distribution indicates enhanced flexibility in the given dihedral.

Let us first consider the critical residues of the RBD region playing vital role in ACE2 interaction. We observe that 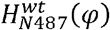 and 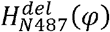 exhibit a sharp peak (fig. 3a), while in rest of the mutated system, increase in flexibility can be found in his degree of freedom due to mutation. The same trend can be found in the backbone dihedral distribution (ψ) of this residue. Sharp unimodal distributions have been observed in 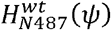 and 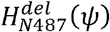, while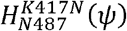, 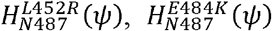 and 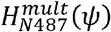 are rather flat (fig. 3b). In this N487, huge increase in flexibility is noticed due to the side chain (χ_1_). The sharp peak of 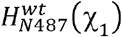 changes into multimodal peaks due to mutations (fig. 3c). 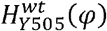 and 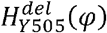 have unimodal distribution, while the mutant variants show relatively flatter distribution depicting enhanced flexibility at that region (fig. 3d). In Y505, 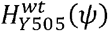 shows a sharp peak, while in rest of the mutated system a multimodal distribution is observed, showing increase in flexibility at this region (fig. 3e). The side chain dihedral (χ_1_) does not show much difference due to mutation (fig. 3f). The cases of the dihedral angles of the other residues forming interacting with host factor of the receptor binding motif are shown in SI figs S1-S6. Overall, the distribution of dihedral angles shows an increase in flexibility in the RBD region due to spike protein mutation.

**Fig3:**
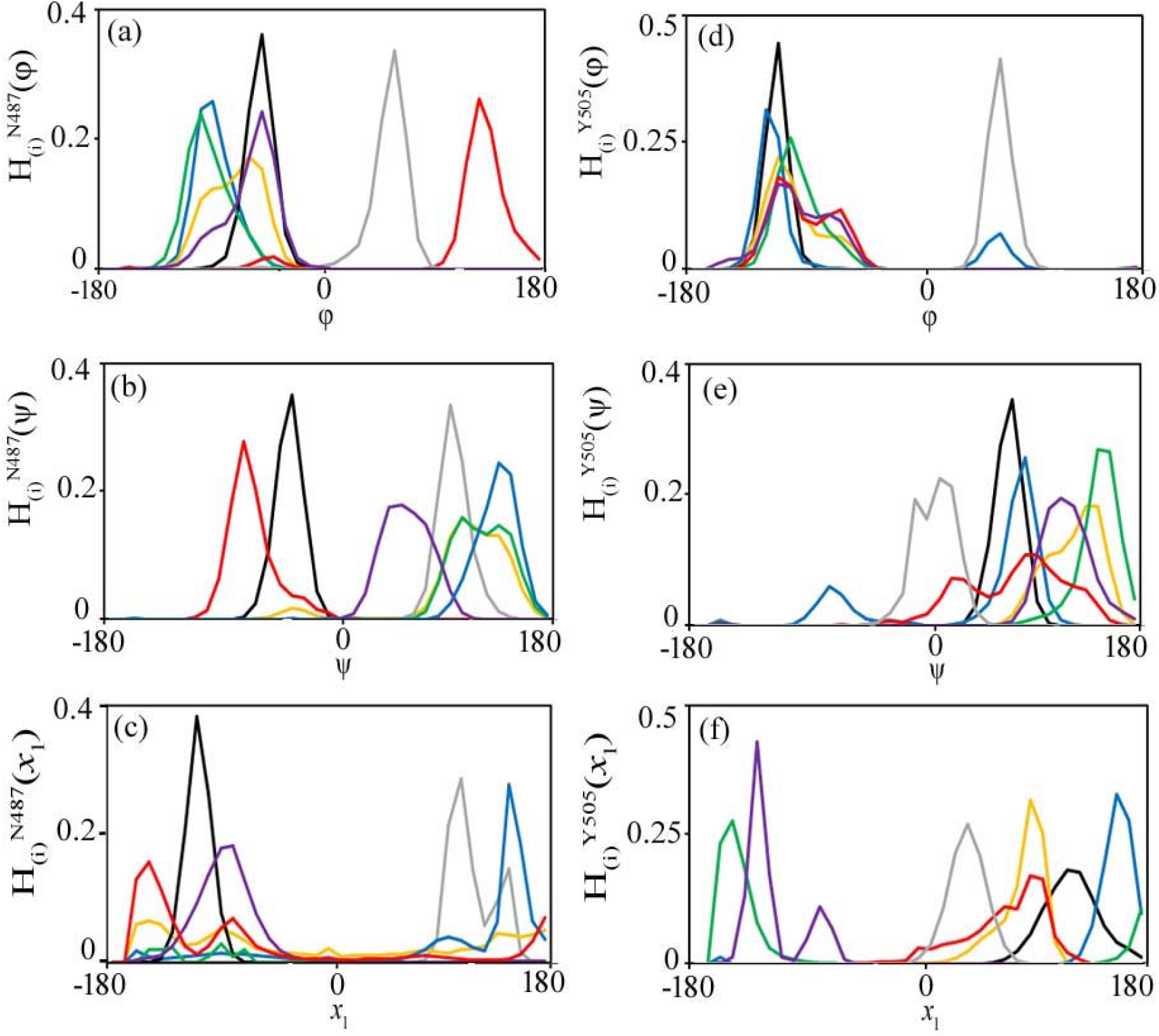
Color online: Histogram distribution of dihedral angles of wild type and variant system at RBD. ACE2 binding motif indicating maximum perturbation is shown here. The color definitions are same as Fig 2. (a) 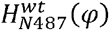 and 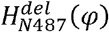 exhibit a sharp peak than the rest of the mutated system, increase in flexibility can be found at this degree of freedom due to mutation. (b) Sharp unimodal distribution have been observed in 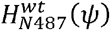 and 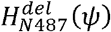. Increase in flexibility is observed due to the flat curves like 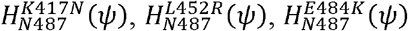, and 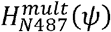. (c) Enhance in flexibility can be noticed due to the side chain (χ_1_). The sharp peak of 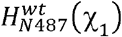 changes into multimodal flattened peak at this degree of freedom due to mutation. (d) 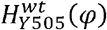 and 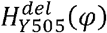 elicits a sharp unimodal distribution and the mutant variants represent flat curved distribution conferring increase in flexibility at this region. (e) 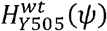 show sharp peak while in rest of the mutated system elicits a multimodal distribution responsible for increase in flexibility. (f) Y505 (χ_1_) does not show much remarkable difference due to mutation.

Let us now consider the dihedral angle distributions of certain residues in the NTD domain. It is noticed that the dihedral distribution *φ* of Y170 is similar for all the system (fig. 4a). 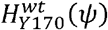 show a sharp peak than the other systems which depicts enhanced flexibility due to mutation (fig. 4b). 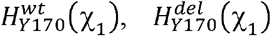, and 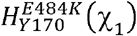 are sharp unimodal, whereas 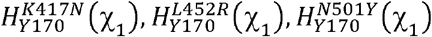 and 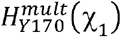 show bimodal distribution (fig. 4c), suggesting increase in flexibility due to such mutations. The cases of the dihedral angles of certain other residues from the NTD domain are shown in SI fig. S7

**Fig4:**
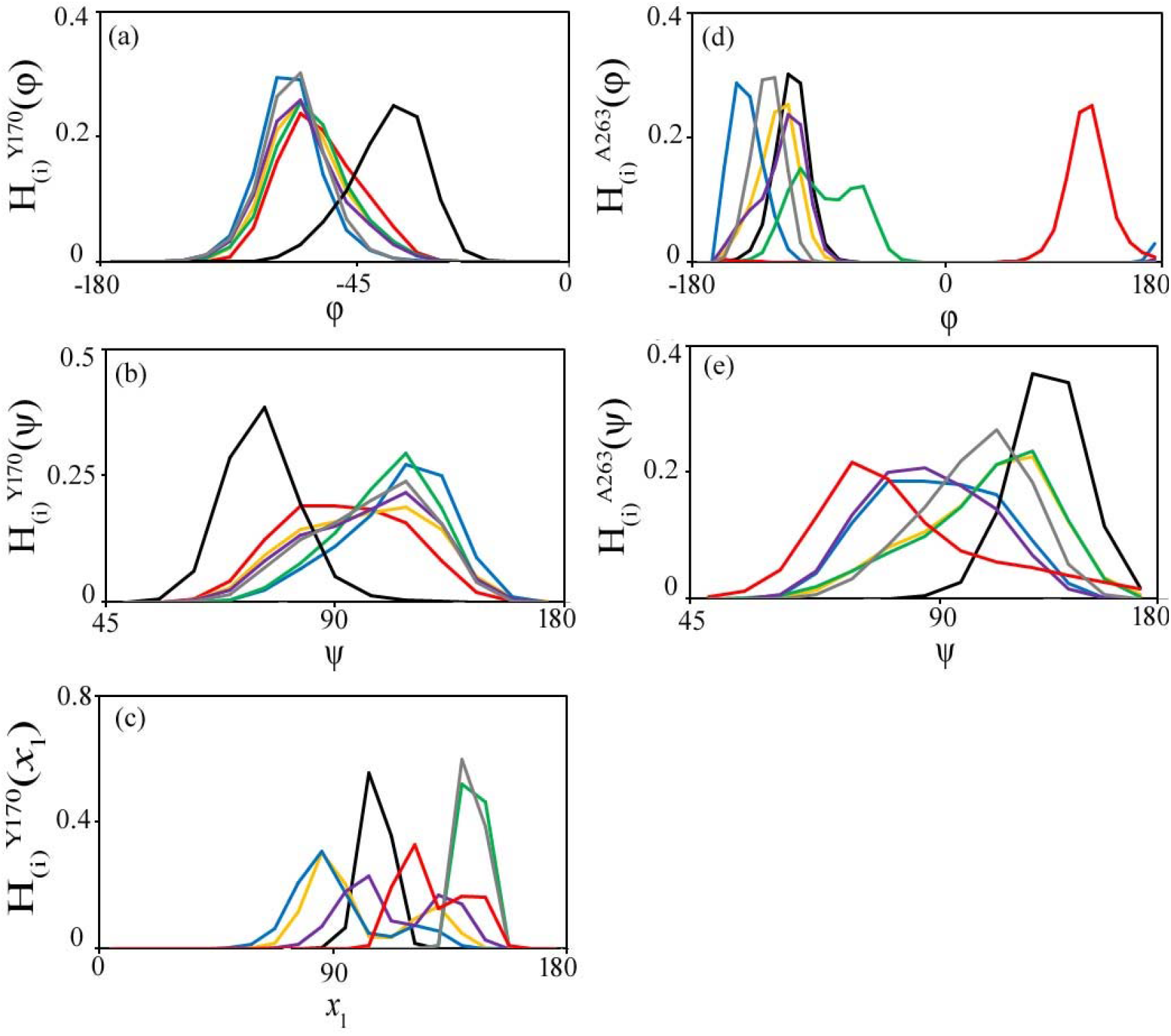
Color online: The dihedral distribution at NTD domain (a) (*φ*)distribution in Y170 is almost uniform for all the system. (b) 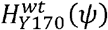 shows a sharp peak than the rest of the system, which depicts elevation of flexibility due to mutation in rest of the system. (c) 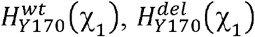 and 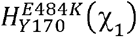 elicit sharp unimodal peak whereas 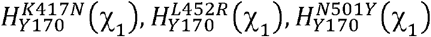 and 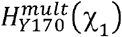 show bimodal distribution. (d) 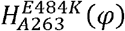 is bimodalwhile, the rest of the system show almost uniform unimodal distribution. (e) 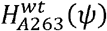 is sharper than the rest of the system.

Now we consider residues from the linker loop. It is observed that 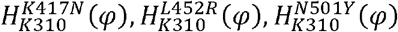 and 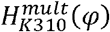 have sharp unimodal peaks, while 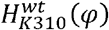 and 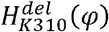 show increase in flexibility (fig. 5a). Thus, mutation causes decrease in flexibility in this degree of freedom. Similarly, for ≠ of K310, decrease in flexibility is observed particularly in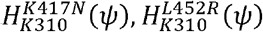 and 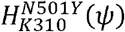(fig. 5b). The side chain dihedral (χ_1_) of K310 shows more flexibility in 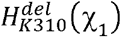 and 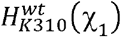 than 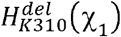 and 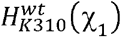. (fig. 5c). 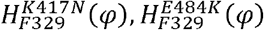 and 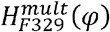 exhibit a sharp tall unimodal peak suggesting that the flexibility is lost after mutation (fig. 3d). The sharp peak of 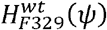 depicts decrease in flexibility due to such mutation, whereas in rest of the system the increase in flexibility is prominent (fig. 3e). In the side chain dihedral (χ_1_) of F329, the unimodal acute peak in 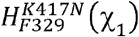 becomes compressed and bimodal distribution in different other systems illustrating expansion of flexibility at this region (fig. 3f). Dihedral distributions of rest of the residues from the region are shown in SI Fig S8-S10.

**Fig5:**
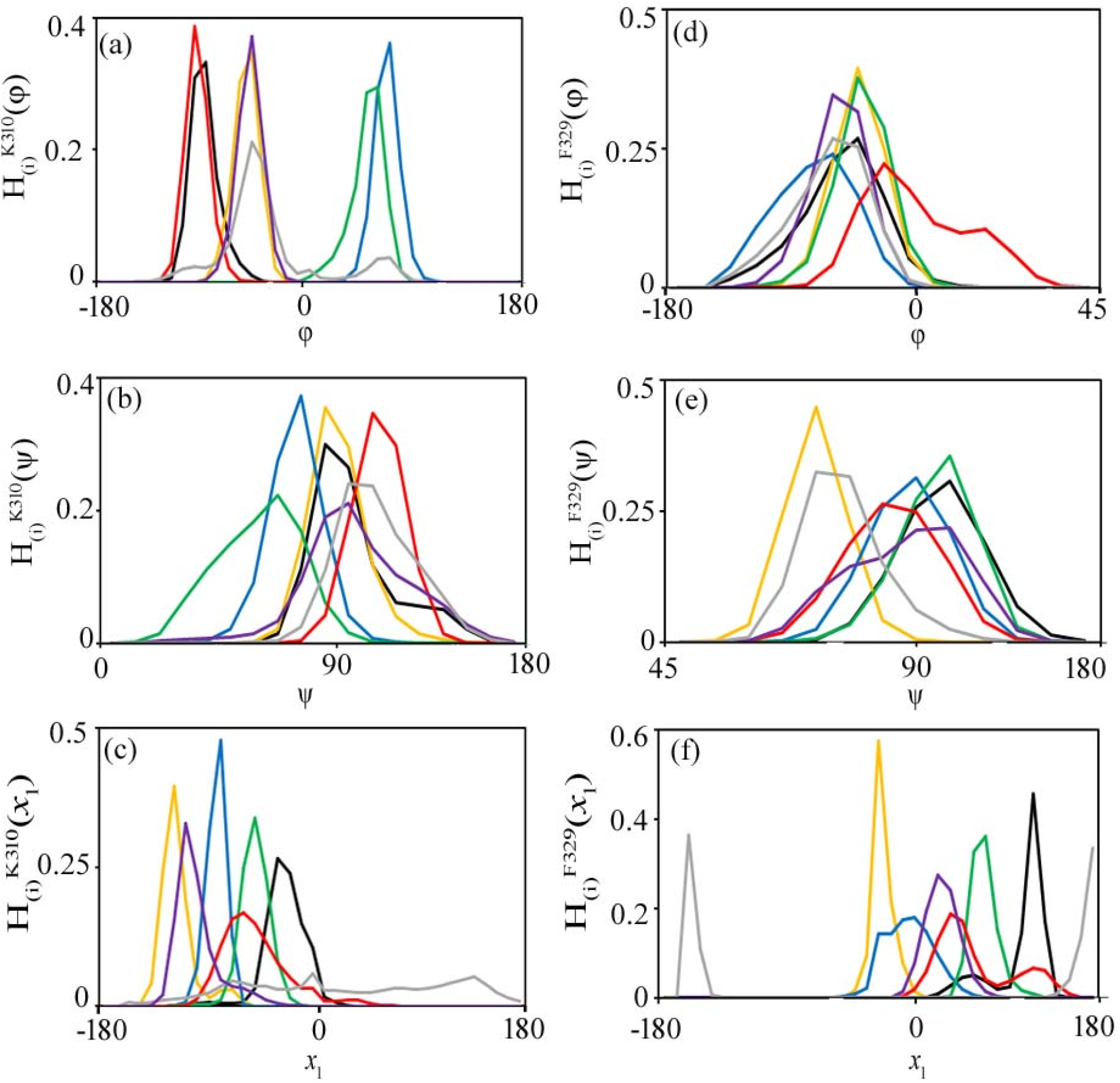
Color online: Histogram distribution of dihedral angles of wild type and variant system at linker loop. The color representations are followed from Fig2. (a) *Increase* in flexibility is observed in 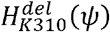(b) Decrease in flexibility can be observed in 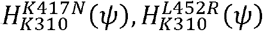 and 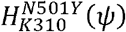(c) Rigidity in the dihedral angle distribution due to side chain (χ_1_) is conferred in most of the mutant variants. (d)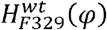 and 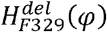 exhibit a bell-shaped flattened peak than the tall sharper peak of the mutated systems. (e) The sharp peak of 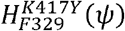 illustrates the decrease in flexibility over 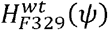. Distribution in rest of the system are almost uniform. (f) In the side chain dihedral (χ_1_) of F329, the unimodal acute peak in 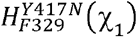 depicts maximum rigidity due to this degree of freedom.

We account for changes in free energy ΔG_i_^conf^ and entropy TΔS_i_^conf^ of conformational changes of the mutated systems with respect to wt-spike from the distributions of the dihedral angles. Positive values of the changes in free energy and entropy indicate destabilization and disorder of the mutated system with respect to the wt-spike, while the negative values indicate stabilization and order in the mutated system. The overall changes in conformational thermodynamics of the various domains of spike protein are obtained by adding all the dihedral contributions from the residues of the particular region. We observe that (Table 1) NTD becomes energetically destabilized and disordered in the mutant variants than wt-spike system. However, NTD from the mt(del)-spike remain stabilized and more ordered with respect to wt-spike. The instability and disorder in the NTD arm are primarily dominated by backbone fluctuations (table 1). In mt(del)-spike, the side chain dihedral imparts maximum stability and ordering. The RBD residues of the spike protein undergo disorder and destabilization in the mutated system compared to the wt-spike system (Table 2). We observe that the mt(del)-spike system remains energetically stabilized and ordered with reference to wt-spike system. This trend is similar as found for NTD arm of the different systems of spike protein. The major contributions in free energy and entropy changes of the RBD region come from backbone dihedrals. It has been found that the overall entropy and free energy changes of the linker loop remains marginal (Table 3), although the residues show more order and stabilization in the mutated case. This suggests that the linker residues play a role like a hinge to control the opening between RBD and NTD. It may be noted that the mt(del)-spike system, the linker shows disorder and destabilization where no hinge role is needed.

**Table 1:** Comparative changes in the conformational thermodynamics (KJ/mol) of the N-terminal region of spike protein in the different mutated systems to wildtype.

| System | $T\Delta S_i^{\text{conf}}$ | | | | $\Delta G_i^{\text{conf}}$ | | | |
| --- | --- | --- | --- | --- | --- | --- | --- | --- |
| | □ | $\psi$ | $\chi_1$ | Total | □ | $\psi$ | $\chi_1$ | Total |
| mt(K417N)-spike | 22.21 | 29.35 | 37.82 | 89.38 | 7.86 | 8.2 | 13.04 | 29.1 |
| mt(L452R)-spike | 13.67 | 29.06 | 19.87 | 62.6 | 9.87 | 8.54 | 5.56 | 7.99 |
| mt(E484K)-spike | 22.11 | 22.18 | 3.56 | 47.85 | 6.47 | 13.93 | -1.12 | 19.28 |
| mt(N501Y)-spike | 4.37 | 4.22 | 13.33 | 21.92 | 7.66 | 5.74 | 6.56 | 19.96 |
| mt(mult)-spike | 24.07 | 28.63 | 19.66 | 72.36 | 21.31 | 10.71 | -4.98 | 27.04 |
| mt(del)-spike | 1.48 | 10.53 | -15.44 | -3.43 | 10.28 | -1.96 | -25.02 | -16.7 |

**Table 2:**
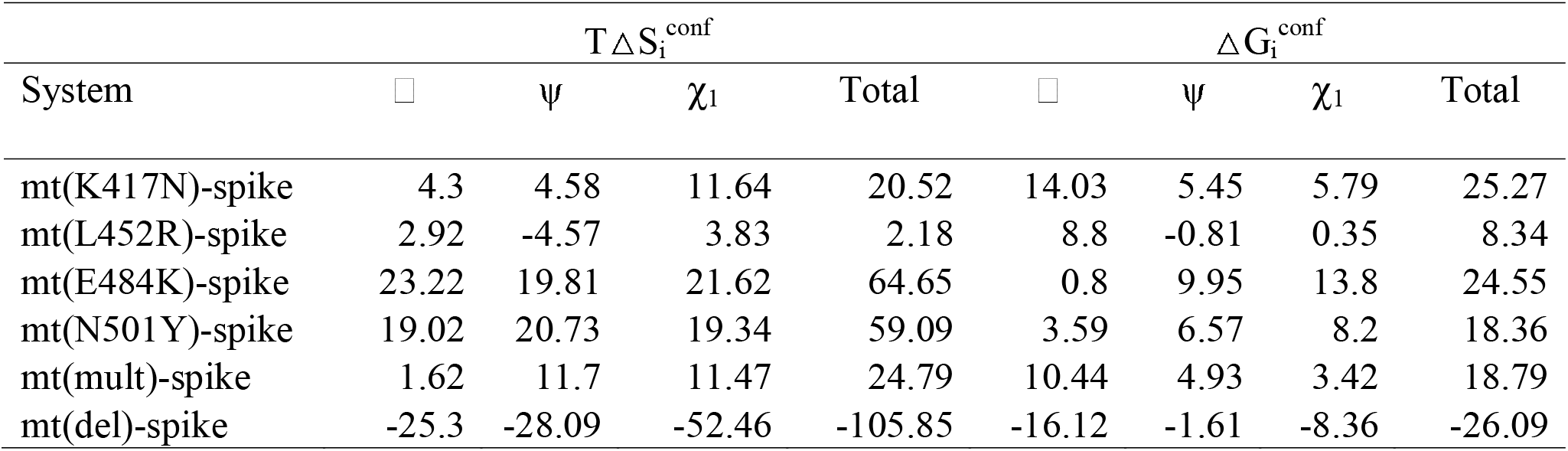
Comparative changes in the conformational thermodynamics (KJ/mol) of the receptor binding domain of spike protein in the different mutated systems to wildtype.

**Table 3:** Comparative changes in the conformational thermodynamics (KJ/mol) of the linker loop of spike protein in the different mutated systems to wildtype.

| System | $T\Delta S_i^{\text{conf}}$ | | | | $\Delta G_i^{\text{conf}}$ | | | |
| --- | --- | --- | --- | --- | --- | --- | --- | --- |
| | □ | $\psi$ | $\chi_1$ | Total | □ | $\psi$ | $\chi_1$ | Total |
| mt(K417N)-spike | 0.6 | -0.64 | -0.41 | -0.45 | 0.36 | -0.28 | -0.2 | -0.12 |
| mt(L452R)-spike | -0.52 | -3.65 | -0.17 | -4.34 | 0.39 | -0.56 | 0.32 | 0.15 |
| mt(E484K)-spike | -1.65 | -2.82 | -0.86 | -5.33 | -2.69 | -0.88 | 3.11 | -0.46 |
| mt(N501Y)-spike | -0.52 | -0.17 | 0.17 | -0.52 | -0.05 | -0.12 | -0.2 | -0.37 |
| mt(mult)-spike | -0.93 | -0.17 | 0.32 | -0.78 | -0.83 | -0.15 | -0.03 | -0.65 |
| mt(del)-spike | 1.43 | 1.28 | 2.89 | 5.6 | 0.32 | 0.37 | 0.42 | 1.11 |

Both the residue-wise and the domain wise free energy and entropy costs for conformational changes in the mutated protein with respect to the wt-type protein in the RBD and the linker regions are shown in SI Tables S1-S5. The overall change in entropy in the residues of RBD of mt(K417N)-spike is 5.62 KJ/mol where the major changes are in the backbone dihedrals. The total change in free energy is 7.28 KJ/mol in which the backbone dihedrals contribute the most (SI Table S1). In case of another point mutation mt(L452R)-spike, the total change in entropy of the region is 11.86 KJ/mol, where the backbone dihedrals together account for most of the changes (SI Table S2). The changes in the stability are marginal in all cases except Y449 having the highest change in stability. The total change in free energy is 4.8 KJ/mol (SI Table S2). Altogether change in entropy of mt(E484K)-spike is 1.63 KJ/mol where the backbone dihedrals are the primary contributor. The backbone dihedral (*φ*) is the key factor for changes in free energy of mt(E484K)-spike in this region, the overall change of free energy being 1.66 KJ/mol (SI Table S3). The total changes in entropy of the region for mt(N501Y)-spike is 8.59 KJ/mol, which is exhibited by the total changes in the backbone dihedrals (SI Table S4). The change in free energy is 5.35 KJ/mol where (*φ*) backbone dihedral acts as pivotal for the changes (SI Table S4).The backbone dihedrals contribute for major changes of the region, the total changes in entropy and free energy in mt(mult)-spike being 9.09 KJ/mol and 6.71 KJ/mol respectively (SI Table S5).The overall changes in entropy and free energy in the mt(del)-spike is −10.55 KJ/mol and −7.03 KJ/mol, where the backbone dihedrals are the key factors for such changes in the system (SI Table S6). The details of conformational thermodynamics data for the certain residues of NTD domain are given in SI Table S7-S12.The dihedrals fluctuations show conformational free energy destabilization. Similar increases are observed in the conformational entropy. The maximum destabilization and disorder are through the backbone fluctuations. mt(del)-spike shows similar thermodynamic changes as the other cases in NTD. The thermodynamic data for certain linker residues responsible for critical conformation changes are shown in SI Tables S13-18. The overall changes in the entropy and free energy in mt(K317N)-spike are −9 KJ/mol and −4.34 KJ/mol, the maximum being for F329 (SI Table S13). The backbone dihedral distribution accounts for maximum changes in this system. In mt(L452R)-spike the entropy change and free energy change is −9.85 KJ/mol and −4.99 KJ/mol respectively (SI Table S14). Such ordering and stability in free energy is governed by backbone dihedral fluctuations (SI Table S14). The total changes in entropy for K310, G311, Y313, F329 and P330 in mt(E484K)-spike is −6.81 KJ/mol, primarily due to the backbone dihedral distributions. Besides, the total changes in free energy are −3.23 KJ/mol due to the backbone fluctuations. We find that P330 imparts slight increase of entropy (SI Table S15). In mt(N501Y)-spike, −6.92 KJ/mol is the total change in entropy and the total change in free energy is −4.7 KJ/mol (SI Table S16). However, the changes in entropy and free energy found due to such residues of linker loop in mt(mult)-spike is marginal, −0.53 KJ/mol and −1.87 KJ/mol respectively (SI Table S17). The mt(del)-spike system shows that the total change in entropy is 14.24 KJ/mol and the change in free energy is 3.47 KJ/mol (SI Table S18).

We map the changes in conformational free energy ΔG_i_^conf^ and entropy TΔS_i_^comf^, of individual residues of the mutated systems with respect to wt-spike. Here, we show free energetically stabilized and ordered residues in green and the destabilized and disordered ones in red. Careful examination of the ACE2 interacting residues of RBD of mt(K417N)-spike shows G446, Y449, N487, Y489, T500 and Y505 impart enhanced disorder, the maximum being in Y489. Increase in order is observed in Q493 and G502 (Fig. 6a). It is found that Y449 and Y505 confers major decrease in stability, whereas, in rest of them the free energy change is marginal. In case of another point mutation mt(L452R)-spike, most of the interface residues which forms crucial interaction with host factor ACE2 gets disordered (Fig. 6b). G446 and Y449 undergo maximum decrease in order. Q493 shows marginal increase in order. Y449 shows maximum destabilization in free energy of the region of mt(L452R)-spike. G446, N487, Y489, Q493, T500 and Y505 of mt(L452R)-spike show slight decrease in stability due to free energy change. G502 of mt(L452R)-spike account for minor increase in stability. The RBD residues of spike protein of mt(E484K)-spike show enhanced disorder in G446, N487, Y489 and T500, Y489 having the maximum disorder. On the other hand, Q493, G502, Y505 exhibit increase in order (fig. 6c). It has been found that Y449, N487, Y489 and T500 is responsible for decrease in stability in mt(E484K)-spike, the highest destabilized residue isY449. On the other hand, G446, Q493, G502 and Y505 is responsible for marginal increase in stability in mt(E484K)-spike. Almost all the residues of the region in mt(N501Y)-spike protein shows increase in order except Q493 and G502 (fig. 6d). Y505 is responsible for the maximum disorder. The changes in stability of these residues are marginal for the mutation N501Y except Y449, which imparts maximum increase in destabilization (fig. 6d). We find enhance in stability and order in Q493 and G502 in mt(N501Y)-spike. In mt(mult)-spike Y449, N487, Y489, G502 and Y505 shows increase in disorder and disability, the maximum disorder being in Y489. Y449 imparts largest increase in destabilization (fig. 6e). G446 and Q493 show only marginal increase in order and stability. All the RBD residues interacting with ACE2 in the mt(del)-spike remain ordered and stabilized except N487. Y489 shows maximum increase in order and Y449 imparts maximum change in stability (fig. 6f).

**Fig. 6:**
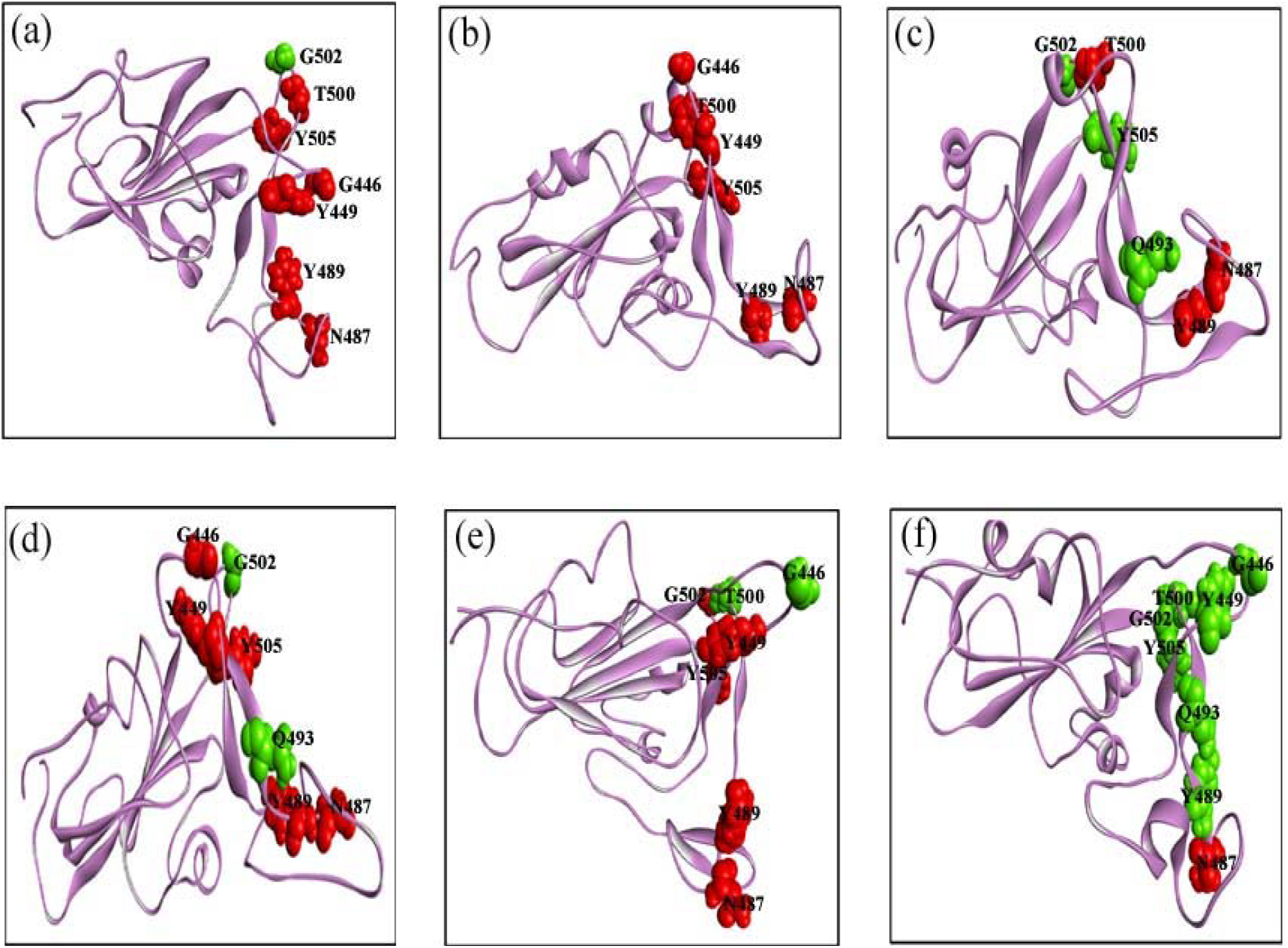
Color online: Illustration of the conformational thermodynamic changes of RBD domain on the average structure of wildtype and mutant variant of spike protein. The destabilized and disordered residues are marked in red (ball-like model). The residues stabilized and ordered are shown in green (ball-like model). The crucial residues forming vital interaction with host factor are only highlighted: (a) mt(K417N)-spike (b) mt(L452R)-spike (c) mt(E484K)-spike (d) mt(N501Y)-spike (e) mt(mult)-spike (f) mt(del)-spike.

Let us now consider the case of linker residues. In mt(K317N)-spike all the residues are ordered and stabilized (fig. 7a). In mt(L452R)-spike K310, G311, Y313, F329 and P330 are ordered as well as stabilized (fig. 7b). K310 imparts maximum increase in order and stability. Fig. 7c illustrates the five linker loop residues of mt(E484K)-spike which undergo ordering and stabilization compared to wt-spike, where K310 accounts for largest change in entropy. The same five residues from the linker loop in mt(N501Y)-spike remains ordered and stabilized with respect to wt-spike (fig. 7d) where G311 of mt(N501Y)-spike account for maximum changes. We observe that the dihedral distribution of K310, G311, Y313, F329 and P330 in mt(mult)-spike are ordered and stabilized (fig. 7e). The residues in mt(del)-spike shows disorder, Y313 being the maximum. Y313, F329 and P330 illustrate instability where F329 accounts for maximum stability, although K310 and G311 show slight increase in stability (fig. 7f). Overall, the dihedral distributions in the linker loop show loss of flexibility in the mutated variants. The decrease in flexibility of the linker region in the mutant variants acts as a hinge to maintain articulation of the opening and closing movement between NTD and RBD. It has been found that mt(del)-spike preferred to attain the close conformation in the equilibrated state.

**Fig. 7:**
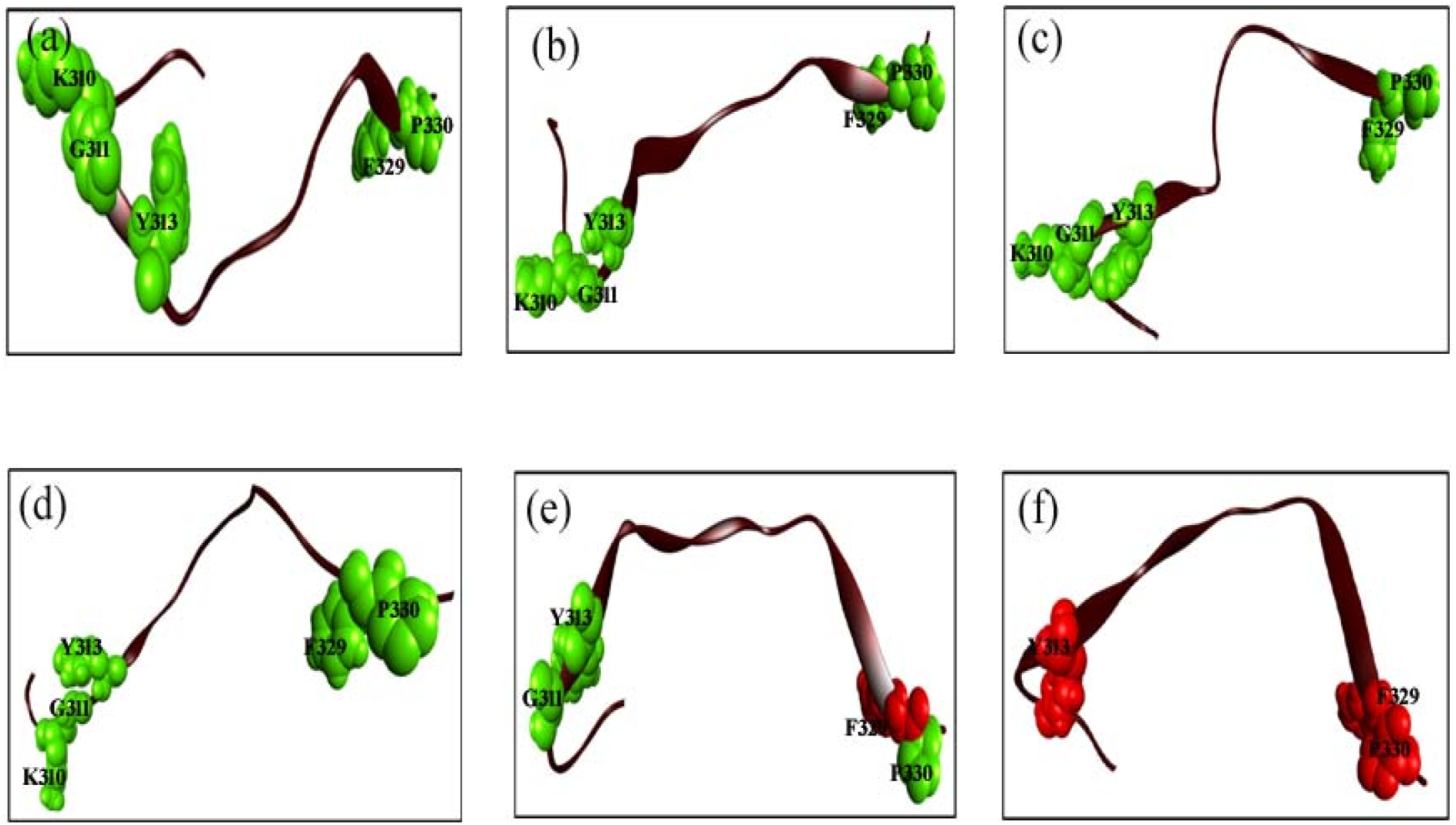
Color online: Close view of the conformational thermodynamic changes of linker region acting as a hinge in spike protein conformation state variation on the average structure of wildtype and mutant variant of spike protein The destabilized and disordered residues are marked in red (ball-like model). The residues stabilized and ordered are shown in green (ball-like model). The residues of the linker responsible for maximum perturbation are shown here: (a) mt(K417N)-spike (b) mt(L452R)-spike (c) mt(E484K)-spike (d) mt(N501Y)-spike (e) mt(mult)-spike (f) mt(del)-spike.

To study the effect of the mutations on ACE2 recognition, we dock the native spike protein wt-spike and each of the mutated spike protein mt(del)-spike with ACE2 where the disordered and destabilized residues are taken to be active residues^30^. The docked complexes are shown in SI Fig. S11. The docking results are shown in SI Table S19. The binding free energy ΔG and the dissociation constant (K_d_)at 25°C are reported from the docking studies. Upon complexed with ACE2, the mutant variants of spike protein show comparatively better interaction than wt-spike and mt(del)-spike. The order of binding affinity of ACE2 is as follows: in mt(E484K)-spike > mt(L452R)-spike > mt(mult)-spike > mt(N501Y)-spike > mt(K417N)-spike compared to that in wt-spike (ΔG=-9.4Kcal/mol) and mt(del)-spike (ΔG=-9.0Kcal/mol) cases. The number of interactions at RBD-ACE2 interface in different system is shown in SI Table S20. The mutant RBD forms more interfacial hydrogen bonds imparting better binding with ACE2. While the stable and ordered linker residues help in stabilizing the open conformation in the mutated protein, the instability and disorder in the RBD facilitates the binding to ACE2.

## 5. Conclusions

In conclusion, we have performed detailed in-silico analysis of stability and order of the spike protein of SARS-CoV-2, primarily responsible for interaction with human cell receptor ACE2. We find open conformation of the mutated protein, while a closed conformation of the wild type protein. The RBD shows instability and disordered residues, while the linker region shows stability and order under mutation compared to the wild-type variant. This may help the ACE2 binding that leads to higher infectivity of the mutated species.

## Author contributions

AMG curate analyzed and interpreted the data and wrote the manuscript. JC interpreted the data, reviewed and edited the manuscript.

## Acknowledgements

AMG is thankful to the Technology Research Centre, S.N. Bose National Centre for Basic Sciences, Kolkata for the computational facilities and the Council for Scientific and Industrial Research for financial support through Research Associateship.

