## Supplementary tables and figures for "Effect on the conformations of spike protein of SARS-CoV-2 due to mutation"

**Author:** Aayatti Mallick Gupta^1*^and Jaydeb Chakrabarti^1^

**Affiliations:**

^1^ Department of Chemical, Biological & Macro-Molecular Sciences, S. N. Bose National Centre for Basic Sciences, Block-JD, Sector-III, Salt Lake, Kolkata-700 106

* To whom correspondence should be addressed:

Corresponding Author: Aayatti Mallick Gupta

Address: Department of Chemical, Biological & Macro-Molecular Sciences, S. N. Bose National Centre for Basic Sciences, Block-JD, Sector-III, Salt Lake, Kolkata-700 106

Supplementary table

Table S1: The changes in conformational thermodynamics (KJ/mol) in the residues of RBM of mt(K417N)-spike with respect to wt-spike

|  | T△S_i_^conf^ | | | | △G_i_^conf^ | | | |
| --- | --- | --- | --- | --- | --- | --- | --- | --- |
| Residue | ϕ | ψ | χ_1_ | Total | ϕ | ψ | χ_1_ | Total |
| G446 | -0.57 | 0.85 | NA | 0.28 | -0.07 | 0.1 | NA | 0.03 |
| Y449 | 0.02 | 0.88 | -0.3 | 0.6 | 3.9 | 0.12 | 0.07 | 4.09 |
| N487 | 0.5 | 0.93 | 0.2 | 1.63 | 0.42 | 0.3 | 0.22 | 0.94 |
| Y489 | 0.12 | 0.42 | 1.89 | 2.43 | 0.05 | 0.12 | 0.1 | 0.27 |
| Q493 | 0.25 | 0.63 | -1.28 | -0.4 | 0.25 | -0.02 | 0.42 | 0.65 |
| T500 | -0.25 | -0.02 | 1.05 | 0.78 | 0.07 | 0.05 | 0.17 | 0.29 |
| G502 | -0.88 | -1.15 | NA | -2.03 | -0.12 | -0.25 | NA | -0.37 |
| Y505 | 1.33 | 0.85 | 0.15 | 2.33 | 0.7 | 0.7 | -0.02 | 1.38 |
| Sum | 0.52 | 3.39 | 1.71 | 5.62 | 5.2 | 1.12 | 0.96 | 7.28 |

Table S2: The changes in conformational thermodynamics (KJ/mol) in the residues of RBM of mt(L452R)-spike with respect to wt-spike

|  | T△S_i_^conf^ | | | | △G_i_^conf^ | | | |
| --- | --- | --- | --- | --- | --- | --- | --- | --- |
| Residue | ϕ | ψ | χ_1_ | Total | ϕ | ψ | χ_1_ | Total |
| G446 | 2.09 | 1.31 | NA | 3.4 | 0.25 | 0.07 | NA | 0.32 |
| Y449 | 0.3 | 0.65 | 2.16 | 3.11 | 3.12 | -0.05 | -0.52 | 2.55 |
| N487 | -0.02 | 0.37 | 0.55 | 0.9 | -0.02 | 0.27 | 0.2 | 0.45 |
| Y489 | 0.27 | 0.02 | 0.52 | 0.81 | 0.52 | -0.05 | 0.32 | 0.79 |
| Q493 | -0.15 | -0.27 | 0.37 | -0.05 | -0.07 | 0.12 | 0.02 | 0.07 |
| T500 | 0.45 | 0.1 | 0.63 | 1.18 | -0.17 | 0.05 | 0.3 | 0.18 |
| G502 | -0.6 | 0.78 | NA | 0.18 | -0.07 | -0.17 | NA | -0.24 |
| Y505 | 0.73 | 0.4 | 1.2 | 2.33 | 0.68 | 0.02 | -0.02 | 0.68 |
| Sum | 3.07 | 3.36 | 5.43 | 11.86 | 4.24 | 0.26 | 0.3 | 4.8 |

Table S3: The changes in conformational thermodynamics (KJ/mol) in the residues of RBM of mt(E484K)-spike with respect to wt-spike

|  | T△S_i_^conf^ | | | | △G_i_^conf^ | | | |
| --- | --- | --- | --- | --- | --- | --- | --- | --- |
| Residue | ϕ | ψ | χ_1_ | Total | ϕ | ψ | χ_1_ | Total |
| G446 | 1.33 | 1.15 | NA | 2.48 | -0.17 | -0.88 | NA | -1.05 |
| Y449 | -1 | -0.02 | 0.6 | -0.42 | 3.22 | -0.02 | 0.05 | 3.25 |
| N487 | 0.95 | 1.26 | -0.05 | 2.16 | 1.03 | 0.17 | -0.55 | 0.65 |
| Y489 | 0.75 | 1 | 1.63 | 3.38 | 0.7 | 0.78 | 0.02 | 1.5 |
| Q493 | 0.02 | 0.02 | -2.41 | -2.37 | -0.05 | -0.85 | -0.45 | -1.35 |
| T500 | -0.27 | -0.63 | 1.36 | 0.46 | -0.07 | -0.12 | 0.25 | 0.06 |
| G502 | -1.31 | -1.2 | NA | -2.51 | -0.27 | -0.25 | NA | -0.52 |
| Y505 | 0.15 | -0.42 | -1.28 | -1.55 | 0.12 | 0.2 | -1.2 | -0.88 |
| Sum | 0.62 | 1.16 | -0.15 | 1.63 | 4.51 | -0.97 | -1.88 | 1.66 |

Table S4: The changes in conformational thermodynamics (KJ/mol) in the residues of RBM of mt(N501Y)spike with respect to wt-spike

|  | T△S_i_^conf^ | | | | △G_i_^conf^ | | | |
| --- | --- | --- | --- | --- | --- | --- | --- | --- |
| Residue | ϕ | ψ | χ_1_ | Total | ϕ | ψ | χ_1_ | Total |
| G446 | 2.09 | 1.42 | NA | 3.51 | 1.56 | 0.22 | NA | 1.78 |
| Y449 | 0.69 | 0.08 | 0.52 | 1.29 | 3.11 | -0.14 | -0.12 | 2.85 |
| N487 | 0.24 | 0.06 | 0.32 | 0.62 | 0.15 | 0.02 | 0.1 | 0.27 |
| Y489 | -0.02 | 0.54 | 1.1 | 1.62 | 0.1 | 0.31 | 0.97 | 1.38 |
| Q493 | -0.19 | 0.36 | -2.01 | -1.84 | -0.08 | -0.27 | -0.85 | -1.2 |
| T500 | -1.13 | 0.14 | 1.49 | 0.5 | -0.15 | -0.02 | 0.11 | -0.06 |
| G502 | -0.66 | -0.83 | NA | -1.49 | -0.12 | -0.2 | NA | -0.32 |
| Y505 | 1.43 | 1.94 | 1.01 | 4.38 | 0.27 | 0.27 | 0.11 | 0.65 |
| Sum | 2.45 | 3.71 | 2.43 | 8.59 | 4.84 | 0.19 | 0.32 | 5.35 |

Table S5: The changes in conformational thermodynamics (KJ/mol) in the residues of RBM of mt(mult)-spike with respect to wt-spike

|  | T△S_i_^conf^ | | | | △G_i_^conf^ | | | |
| --- | --- | --- | --- | --- | --- | --- | --- | --- |
| Residue | ϕ | ψ | χ_1_ | Total | ϕ | ψ | χ_1_ | Total |
| G446 | -0.58 | -0.03 | NA | -0.61 | -0.04 | 0.02 | NA | -0.02 |
| Y449 | -0.02 | 1.26 | -0.04 | 1.2 | 3.28 | 0.11 | 0.06 | 3.45 |
| N487 | 0.59 | 0.9 | 1.37 | 2.86 | 0.73 | 0.31 | 0.22 | 1.26 |
| Y489 | 0.48 | 0.69 | 1.98 | 3.15 | 0.18 | 0.2 | -0.01 | 0.37 |
| Q493 | 0.73 | 1.17 | -2.43 | -0.53 | 0.68 | -0.04 | -0.24 | 0.4 |
| T500 | -0.9 | -0.45 | 0.45 | -0.9 | -0.25 | -0.12 | 0.14 | -0.23 |
| G502 | 1.33 | 1.33 | NA | 2.66 | 0.08 | 0.31 | NA | 0.39 |
| Y505 | 1.27 | 0.29 | -0.3 | 1.26 | 0.41 | 0.01 | 0.67 | 1.09 |
| SUM | 2.9 | 5.16 | 1.03 | 9.09 | 5.07 | 0.8 | 0.84 | 6.71 |

Table S6: The changes in conformational thermodynamics (KJ/mol) in the residues of RBM of mt(del)-spike with respect to wt-spike

|  | T△S_i_^conf^ | | | | △G_i_^conf^ | | | |
| --- | --- | --- | --- | --- | --- | --- | --- | --- |
| Residue | ϕ | ψ | χ_1_ | Total | ϕ | ψ | χ_1_ | Total |
| G446 | -1.47 | -0.95 | NA | -2.42 | -1.32 | -0.1 | NA | -1.42 |
| Y449 | -0.02 | -0.58 | -0.09 | -0.69 | -3.38 | -0.04 | 0.12 | -3.3 |
| N487 | 0.56 | 1.18 | 1.67 | 3.41 | 0.24 | 0.19 | 0.35 | 0.78 |
| Y489 | -0.01 | -1.8 | -2.82 | -4.63 | -0.16 | -0.53 | -0.05 | -0.74 |
| Q493 | -0.42 | -1.04 | -0.8 | -2.26 | -0.01 | -0.11 | -0.89 | -1.01 |
| T500 | -0.02 | 0.93 | -1.14 | -0.23 | -0.12 | 0.05 | -0.24 | -0.31 |
| G502 | -1.81 | -1.11 | NA | -2.92 | -0.3 | -0.21 | NA | -0.51 |
| Y505 | -0.21 | -0.67 | 0.07 | -0.81 | 0.19 | 0.08 | -0.79 | -0.52 |
| SUM | -3.4 | -4.04 | -3.11 | -10.55 | -4.86 | -0.67 | -1.5 | -7.03 |

Table S7: The changes in conformational thermodynamics (KJ/mol) in the residues of NTD of mt(K417N)spike with respect to wt-spike

|  | T△S_i_^conf^ | | | | △G_i_^conf^ | | | |
| --- | --- | --- | --- | --- | --- | --- | --- | --- |
| Residue | ϕ | ψ | χ_1_ | Total | ϕ | ψ | χ_1_ | Total |
| Y170 | 0.01 | 0.09 | 2.38 | 2.48 | -0.25 | 3.22 | 2 | 4.97 |
| A263 | 0.46 | -0.19 | NA | 0.27 | 0.43 | 0.42 | NA | 0.85 |
| Sum | 0.47 | -0.1 | 2.38 | 2.75 | 0.18 | 3.64 | 2 | 5.82 |

Table S8: The changes in conformational thermodynamics (KJ/mol) in the residues of NTD of mt(L452R)spike with respect to wt-spike

|  | T△S_i_^conf^ | | | | △G_i_^conf^ | | | |
| --- | --- | --- | --- | --- | --- | --- | --- | --- |
| Residue | ϕ | ψ | χ_1_ | Total | ϕ | ψ | χ_1_ | Total |
| Y170 | -0.48 | -0.45 | 2.41 | 1.48 | -0.4 | 2.65 | 1.9 | 4.15 |
| A263 | -0.15 | -2.06 | NA | -2.21 | 0.19 | 3.17 | NA | 3.36 |
| Sum | -0.63 | -2.51 | 2.41 | -0.73 | -0.21 | 5.82 | 1.9 | 7.51 |

Table S9: The changes in conformational thermodynamics (KJ/mol) in the residues of NTD of mt(E484K)spike with respect to wt-spike

|  | T△S_i_^conf^ | | | | △G_i_^conf^ | | | |
| --- | --- | --- | --- | --- | --- | --- | --- | --- |
| Residue | ϕ | ψ | χ_1_ | Total | ϕ | ψ | χ_1_ | Total |
| Y170 | 0.01 | -0.48 | -0.62 | -1.09 | -0.24 | -3.29 | 0.09 | -3.44 |
| A263 | 1.56 | -0.19 | NA | 1.37 | 1.9 | 4.09 | NA | 5.99 |
| Sum | 1.57 | -0.67 | -0.62 | 0.28 | 1.66 | 0.8 | 0.09 | 2.55 |

Table S10: The changes in conformational thermodynamics (KJ/mol) in the residues of NTD of mt(N501Y)spike with respect to wt-spike

|  | T△S_i_^conf^ | | | | △G_i_^conf^ | | | |
| --- | --- | --- | --- | --- | --- | --- | --- | --- |
| Residue | ϕ | ψ | χ_1_ | Total | ϕ | ψ | χ_1_ | Total |
| Y170 | -0.16 | -0.4 | 1.53 | 0.97 | -0.34 | 3 | 0.19 | 2.85 |
| A263 | 0.48 | -2.34 | NA | -1.86 | 0.22 | 3.6 | NA | 3.82 |
| Sum | 0.32 | -2.74 | 1.53 | -0.89 | -0.12 | 6.6 | 0.19 | 6.67 |

Table S11: The changes in conformational thermodynamics (KJ/mol) in the residues of NTD of mt(mult)spike with respect to wt-spike

|  | T△S_i_^conf^ | | | | △G_i_^conf^ | | | |
| --- | --- | --- | --- | --- | --- | --- | --- | --- |
| Residue | ϕ | ψ | χ_1_ | Total | ϕ | ψ | χ_1_ | Total |
| Y170 | -0.09 | 0.02 | 2.59 | 2.52 | -0.2 | 3.22 | 0.53 | 3.55 |
| A263 | 0.83 | -0.08 | NA | 0.75 | 0.49 | 3.2 | NA | 3.69 |
| Sum | 0.74 | -0.06 | 2.59 | 3.27 | 0.29 | 6.42 | 0.53 | 7.24 |

Table S12: The changes in conformational thermodynamics (KJ/mol) in the residues of NTD of mt(del)spike with respect to wt-spike

|  | T△S_i_^conf^ | | | | △G_i_^conf^ | | | |
| --- | --- | --- | --- | --- | --- | --- | --- | --- |
| Residue | ϕ | ψ | χ_1_ | Total | ϕ | ψ | χ_1_ | Total |
| Y170 | 0.89 | 1.45 | 4.05 | 6.39 | 0.25 | 0.56 | 0.93 | 1.74 |
| A263 | 1.31 | 1.49 | NA | 2.8 | 0.34 | 0.49 | NA | 0.83 |
| Sum | 2.2 | 2.94 | 4.05 | 9.19 | 0.59 | 1.05 | 0.93 | 2.57 |

Table S13: The changes in conformational thermodynamics (KJ/mol) in the residues of linker of mt(K417N)spike with respect to wt-spike

|  | T△S_i_^conf^ | | | | △G_i_^conf^ | | | |
| --- | --- | --- | --- | --- | --- | --- | --- | --- |
| Residue | ϕ | ψ | χ_1_ | Total | ϕ | ψ | χ_1_ | Total |
| K310 | -0.38 | -0.93 | -0.38 | -1.69 | -0.01 | -0.32 | -0.08 | -0.41 |
| G311 | -1.13 | -0.9 | NA | -2.03 | -0.31 | 0.01 | NA | -0.3 |
| Y313 | 0.16 | 0.21 | -0.63 | -0.26 | 0.27 | -0.26 | -0.41 | -0.4 |
| F329 | -0.99 | -1.26 | -0.71 | -2.96 | -0.54 | -0.76 | -0.75 | -2.05 |
| P330 | -0.52 | -0.52 | -1.02 | -2.06 | -0.41 | -0.71 | -0.06 | -1.18 |
| Sum | -2.86 | -3.4 | -2.74 | -9 | -1 | -2.04 | -1.3 | -4.34 |

Table S14: The changes in conformational thermodynamics (KJ/mol) in the residues of linker of mt(L452R)spike with respect to wt-spike

|  | T△S_i_^conf^ | | | | △G_i_^conf^ | | | |
| --- | --- | --- | --- | --- | --- | --- | --- | --- |
| Residue | ϕ | ψ | χ_1_ | Total | ϕ | ψ | χ_1_ | Total |
| K310 | -0.89 | -1.46 | -1.17 | -3.52 | -0.51 | -0.46 | -0.67 | -1.64 |
| G311 | -1.41 | -1.82 | NA | -3.23 | -1.02 | 0.79 | NA | -0.23 |
| Y313 | 0.01 | -0.08 | -0.52 | -0.59 | -0.01 | -0.69 | -0.4 | -1.1 |
| F329 | -0.51 | -0.79 | 0.38 | -0.92 | -0.26 | -0.3 | 0.02 | -0.54 |
| P330 | -0.41 | 0.38 | -1.56 | -1.59 | -0.27 | -0.59 | -0.62 | -1.48 |
| Sum | -3.21 | -3.77 | -2.87 | -9.85 | -2.07 | -1.25 | -1.67 | -4.99 |

Table S15: The changes in conformational thermodynamics (KJ/mol) in the residues of linker of mt(E484K)spike with respect to wt-spike

|  | T△S_i_^conf^ | | | | △G_i_^conf^ | | | |
| --- | --- | --- | --- | --- | --- | --- | --- | --- |
| Residue | ϕ | ψ | χ_1_ | Total | ϕ | ψ | χ_1_ | Total |
| K310 | -0.83 | -0.56 | -1.3 | -2.69 | -0.51 | 0.02 | -0.26 | -0.75 |
| G311 | -0.25 | -1.22 | NA | -1.47 | -0.97 | 0.18 | NA | -0.79 |
| Y313 | -0.16 | -0.16 | -0.05 | -0.37 | -0.04 | -0.26 | -0.21 | -0.51 |
| F329 | -0.76 | -0.53 | -0.24 | -1.53 | -0.52 | -0.23 | -0.14 | -0.89 |
| P330 | -0.37 | -0.5 | 0.12 | -0.75 | -0.33 | -0.16 | 0.2 | -0.29 |
| Sum | -2.37 | -2.97 | -1.47 | -6.81 | -2.37 | -0.45 | -0.41 | -3.23 |

Table S16: The changes in conformational thermodynamics (KJ/mol) in the residues of linker of mt(N501Y)spike with respect to wt-spike

|  | T△S_i_^conf^ | | | | △G_i_^conf^ | | | |
| --- | --- | --- | --- | --- | --- | --- | --- | --- |
| Residue | ϕ | ψ | χ_1_ | Total | ϕ | ψ | χ_1_ | Total |
| K310 | -0.69 | -1.34 | 0.12 | -1.91 | -0.36 | -0.34 | -0.05 | -0.75 |
| G311 | -2.55 | -0.93 | NA | -3.48 | -0.99 | -1.4 | NA | -2.39 |
| Y313 | -0.37 | -0.88 | -0.29 | -1.54 | -0.17 | -0.2 | -0.24 | -0.61 |
| F329 | -0.25 | -0.42 | 0.01 | -0.66 | -0.19 | -0.14 | -0.23 | -0.56 |
| P330 | -0.1 | -0.25 | 1.02 | 0.67 | -0.18 | -0.08 | -0.13 | -0.39 |
| Sum | -3.96 | -3.82 | 0.86 | -6.92 | -1.89 | -2.16 | -0.65 | -4.7 |

Table S17: The changes in conformational thermodynamics (KJ/mol) in the residues of linker of mt(mult)-spike with respect to wt-spike

|  | T△S_i_^conf^ | | | | △G_i_^conf^ | | | |
| --- | --- | --- | --- | --- | --- | --- | --- | --- |
| Residue | ϕ | ψ | χ_1_ | Total | ϕ | ψ | χ_1_ | Total |
| K310 | 0.01 | -0.63 | 1.07 | 0.45 | -0.44 | -0.13 | 0.04 | -0.53 |
| G311 | -0.45 | 0.42 | NA | -0.03 | -0.01 | -0.92 | NA | -0.93 |
| Y313 | 0.01 | -0.07 | -0.29 | -0.35 | 0.03 | -0.05 | -0.56 | -0.58 |
| F329 | -0.23 | -0.27 | 0.75 | 0.25 | 0.01 | 0.07 | 0.51 | 0.59 |
| P330 | -0.21 | -0.01 | -0.63 | -0.85 | -0.22 | -0.13 | -0.07 | -0.42 |
| Sum | -0.87 | -0.56 | 0.9 | -0.53 | -0.63 | -1.16 | -0.08 | -1.87 |

Table S18: The changes in conformational thermodynamics (KJ/mol) in the residues of linker of mt(del)-spike with respect to wt-spike

|  | T△S_i_^conf^ | | | | △G_i_^conf^ | | | |
| --- | --- | --- | --- | --- | --- | --- | --- | --- |
| Residue | ϕ | ψ | χ_1_ | Total | ϕ | ψ | χ_1_ | Total |
| K310 | 0.15 | 0.14 | 0.47 | 0.76 | -0.35 | 0.04 | 0.21 | -0.1 |
| G311 | 0.1 | 0.81 | NA | 0.91 | -0.18 | 0.11 | NA | -0.07 |
| Y313 | 1.18 | 0.35 | 3.14 | 4.67 | 0.85 | 0.09 | 0.61 | 1.55 |
| F329 | 0.89 | 1.06 | 1.95 | 3.9 | 0.54 | 0.31 | 0.97 | 1.82 |
| P330 | 1.13 | 0.92 | 1.95 | 4 | 0.17 | -0.42 | 0.52 | 0.27 |
| Sum | 3.45 | 3.28 | 7.51 | 14.24 | 1.03 | 0.13 | 2.31 | 3.47 |

Table S19: Docking results of ACE2 binding due to mutations of RBD of spike protein

| Scores | wt-spike | mt(del) | mt(K417N) | mt(L452R) | mt(E484K) | mt(N501Y) | mt(mult) |
| --- | --- | --- | --- | --- | --- | --- | --- |
| ΔG (kcal mol^-1^) | -9.4 | -9.0 | -10.3 | -11.5 | -12.5 | -11.2 | -11.9 |
| K_d (_at 25°C) | 1.2E-07 | 2.6E-07 | 3.0E-08 | 3.6E-09 | 6.6E-10 | 5.7E-09 | 1.9E-09 |

Table S20: Intermolecular interactions in RBD-ACE2 interface in different mutant variants.

| Complex name | Residue and atom name | Bond distance | Bond type |
| --- | --- | --- | --- |
| wt-spike-Ace2 | **THR27:HG1** - GLN493:OE1 | 2.15547 | Hydrogen Bond |
|  | **ASP30:OD1** - TYR505 | 4.36145 | Electrostatic |
| mt(del)-Ace2 | **TYR83:HH** - ASN501:O | 2.344 | Hydrogen Bond |
|  | **LYS353:HZ2** - GLN493:OE1 | 1.77362 | Hydrogen Bond |
|  | **HIS34** - TYR449 | 4.49915 | Hydrophobic |
| mt(K417N)-Ace2 | ASN487:HD22 - **TYR83:OH** | 2.77532 | Hydrogen Bond |
|  | TYR489:HH - **ASP30:O** | 2.80928 | Hydrogen Bond |
|  | **THR27:HG1** - ASN487:O | 1.99299 | Hydrogen Bond |
|  | **LYS353:HZ3** - GLY446:O | 2.97866 | Hydrogen Bond |
|  | **ASP30:OD2** - TYR489 | 3.92176 | Electrostatic |
|  | TYR489 - **LYS31** | 5.38701 | Hydrophobic |
| mt(L452R)-Ace2 | LYS417:HZ2 - **ASP30:OD2** | 1.6034 | Salt bridge |
|  | TYR489:HH - **TYR83:OH** | 2.03716 | Hydrogen Bond |
|  | VAL503:HN - **LYS353:O** | 1.77912 | Hydrogen Bond |
|  | TYR505:HH - **HIS34:O** | 2.20086 | Hydrogen Bond |
|  | **LYS31:HZ3** - GLN493:OE1 | 1.9584 | Hydrogen Bond |
|  | **THR27:CG2** - TYR489 | 3.83407 | Hydrophobic |
| mt(N501Y)-Ace2 | **LYS353:HZ2** - GLU484:OE2 | 1.69089 | Salt bridge |
|  | **THR27:HG1** - TYR501:O | 1.78589 | Hydrogen Bond |
|  | **LYS31:HZ2** - GLY504:O | 1.8712 | Hydrogen Bond |
|  | **TYR41:HH** - ASN487:OD1 | 2.28748 | Hydrogen Bond |
|  | GLY502:CA - **GLN24:O** | 3.49465 | Hydrogen Bond |
|  | **LYS31:NZ** - TYR505 | 4.38047 | Electrostatic |
|  | TYR505 - **LYS31** | 4.5209 | Hydrophobic |
| mt(E484K-Ace2 | ASN487:HN - **HIS34:NE2** | 2.51421 | Hydrogen Bond |
|  | **HIS34:CE1** - ASN487:OD1 | 3.18126 | Hydrogen Bond |
|  | **ASN330:HD22** - TYR505 | 2.97607 | Hydrogen Bond |
| mt(mult)-Ace2 | THR500:HG1 - **ASP30:OD2** | 1.71191 | Hydrogen Bond |
|  | SER494:CB - **LYS353:O** | 3.41675 | Hydrogen Bond |

_ACE2 residues are marked in bold._

Supplementary figures legends:

Fig. S1: Histogram distribution of dihedral angles of wild type and variant system due to RBD residue G446. The color definitions are wt-spike in black, mt(del)-spike in grey, mt(K417N)-spike in yellow, mt(L452R)-spike in blue, mt(E484K)-spike in green, mt(N501Y)-spike in red and mt(mult)-spike in violet. (a) $H_{G446}^{wt}\left( \varphi\right)$ shows a sharper unimodal peak than the flattened multimodal peak in most of the mutated system depicting increase in flexibility can be found at this degree of freedom due to mutation. (b) Sharp unimodal distribution have been observed in $H_{G446}^{wt}\left( \psi\right)$ and $H_{G446}^{del}\left( \psi\right)$. Increase in flexibility is observed due to the flat multimodal curves like $H_{G446}^{K417N}\left( \psi\right)$, $H_{G446}^{L452R}\left( \psi\right)$, $H_{G446}^{E484K}\left( \psi\right)$ and $H_{G446}^{mult}\left( \psi\right)$.

Fig. S2: Histogram distribution of dihedral angles of wild type and variant system due to RBD residue Y449. The color definitions are same as in fig. S1. (a) $H_{Y449}^{wt}\left( \varphi\right)$ elicits a sharp unimodal distribution and the mutant variants represent flat curved distribution conferring increase in flexibility at this region. (b) $H_{Y449}^{wt}\left( \psi\right)$ and $H_{Y449}^{del}\left( \psi\right)$ show uniform distribution. Changes in flexibility is profound in $H_{Y449}^{K417N}\left( \psi\right)$ and $H_{Y449}^{mult}\left( \psi\right)$. (c) Enhance in flexibility can be noticed due to the side chain (χ_1_). The sharp peak of $H_{Y449}^{wt}\left( \chi_{1} \right)$ changes into multimodal flattened peak at this degree of freedom due to mutation except $H_{Y449}^{L452R}\left( \chi_{1} \right)$.

Fig. S3: Histogram distribution of dihedral angles of wild type and variant system due to RBD residue Y489. The color definitions are same as in fig. S1. (a) Slight increase in flexibility in the dihedral angle distribution is visible at this degree of freedom due to mutation. (b) No such remarkable changes in flexibility can be found due to backbone dihedral $\left( \psi\right).$ (c) $H_{Y489}^{wt}\left( \chi_{1} \right)$ and $H_{Y489}^{del}\left( \chi_{1} \right)$ elicit sharp unimodal peak whereas $H_{Y489}^{K417N}\left( \chi_{1} \right),H_{Y489}^{L452R}\left( \chi_{1} \right),H_{Y489}^{N501Y}\left( \chi_{1} \right),H_{Y489}^{E484K}\left( \chi_{1} \right),$ and $H_{Y170}^{mult}\left( \chi_{1} \right)$ show bimodal distribution.

Fig. S4: Histogram distribution of dihedral angles of wild type and variant system due to RBD residue Q493. The color definitions are same as in fig. S1. (a) $H_{Q493}^{wt}\left( \varphi\right)$ elicits a sharp unimodal distribution and the mutant variants represent some increase in flexibility due to mutation. (b) Increase in flexibility can be found in $H_{Q493}^{K417N}\left( \psi\right)$, $H_{Q493}^{N501Y}\left( \psi\right)$, $H_{Q493}^{E484K}\left( \psi\right)$ and $H_{Q493}^{mult}\left( \psi\right)$. (c) The sharp peak of $H_{Q493}^{wt}\left( \chi_{1} \right)$ and $H_{Q493}^{del}\left( \chi_{1} \right)$changes into multimodal flattened peak at this degree of freedom due to mutation.

Fig. S5: Histogram distribution of dihedral angles of wild type and variant system due to RBD residue T500. The color definitions are same as in fig. S1. (a) The backbone dihedral $\left( \varphi\right)$ shows increase in flexibility in all the variants due to mutation. (b) ) $H_{T500}^{wt}\left( \psi\right)$ elicits a sharp unimodal distribution and the rest of the system represent flat curved distribution conferring increase in flexibility at this region. (c) The sharp spike of $H_{T500}^{wt}\left( \chi_{1} \right)$ and $H_{T500}^{del}\left( \chi_{1} \right)$changes into multimodal flattened peak at this degree of freedom due to mutation.

Fig. S6: Histogram distribution of dihedral angles of wild type and variant system due to RBD residue G502. The color definitions are same as in fig. S1. (a) Enhance in flexibility is remarkable in $\left( \varphi\right)$dihedral due to mutation. (b) The sharp peak of $H_{G502}^{wt}\left( \psi\right)$ changes into multimodal flattened peak in rest of the system.

Fig. S7: Histogram distribution of dihedral angles of wild type and variant system due to NTD residue A263. The color definitions are same as in fig. S1. (a) Increase in flexibility is found in $H_{A263}^{E384}\left( \varphi\right)$ and $H_{A263}^{mult}\left( \varphi\right)$. (b) Flattening of the peak is prominent in $\left( \psi\right)$ due to mutation.

Fig. S8: Histogram distribution of dihedral angles of wild type and variant system due to linker residue G311. The color definitions are same as in fig. S1. (a) The sharp peak of $H_{G311}^{N501Y}\left( \varphi\right)$ becomes multimodal in rest of the system. (b) $H_{G311}^{del}\left( \psi\right)$ becomes multimodal depicting maximum increase in flexibility.

Fig. S9: Histogram distribution of dihedral angles of wild type and variant system due to linker residue Y313. The color definitions are same as in fig. S1. (a) The distribution due to backbone dihedral $\left( \varphi\right)$ does not show much change due to mutation. (b) Decrease in flexibility is remarkable the maximum due to $H_{Y313}^{N501Y}\left( \psi\right)$. $H_{Y313}^{wt}\left( \psi\right)$ and $H_{Y313}^{del}\left( \psi\right)$ show much flexibility due to this degree of freedom. (c) Decrease in flexibility appears profound to the side chain dihedral $\left( \chi_{1} \right)$ due to mutation.

Fig. S10: Histogram distribution of dihedral angles of wild type and variant system due to linker residue P330. The color definitions are same as in fig. S1. (a) The sharpest tapering peak of $H_{P330}^{K417Y}\left( \varphi\right)$ gradually become bell-shaped in $H_{P330}^{del}\left( \varphi\right)$. $H_{P330}^{wt}\left( \varphi\right)$ shows somewhat flattened distribution conferring flexibility of the region is more in wildtype than in the mutant variants. (b) Sharp decrease in flexibity is observed due to mutation except $H_{P330}^{L452R}\left( \psi\right)$. $H_{P330}^{wt}\left( \varphi\right)$ shows bimodal distribution. (c) Flexibity due to side chain dihedral $\left( \chi_{1} \right)$ gets reduced in $H_{P330}^{del}\left( \chi_{1} \right)$and $H_{P330}^{wt}\left( \chi_{1} \right)$.

Fig. S11: Docked complex of spike protein (pink) and ACE2 (green). (a) crystal structure (PDB id:) of co-crystal of RDB region interacting with ACE2. (b) wt-spike-ACE2 (c) mt(K417N)-Ace2 (d) mt(L452R)-Ace2 (e) mt(E484K-Ace2 (f) mt(N501Y)-Ace2 (g) mt(mult)-Ace2 (h) mt(del)-Ace2

Fig. S1:

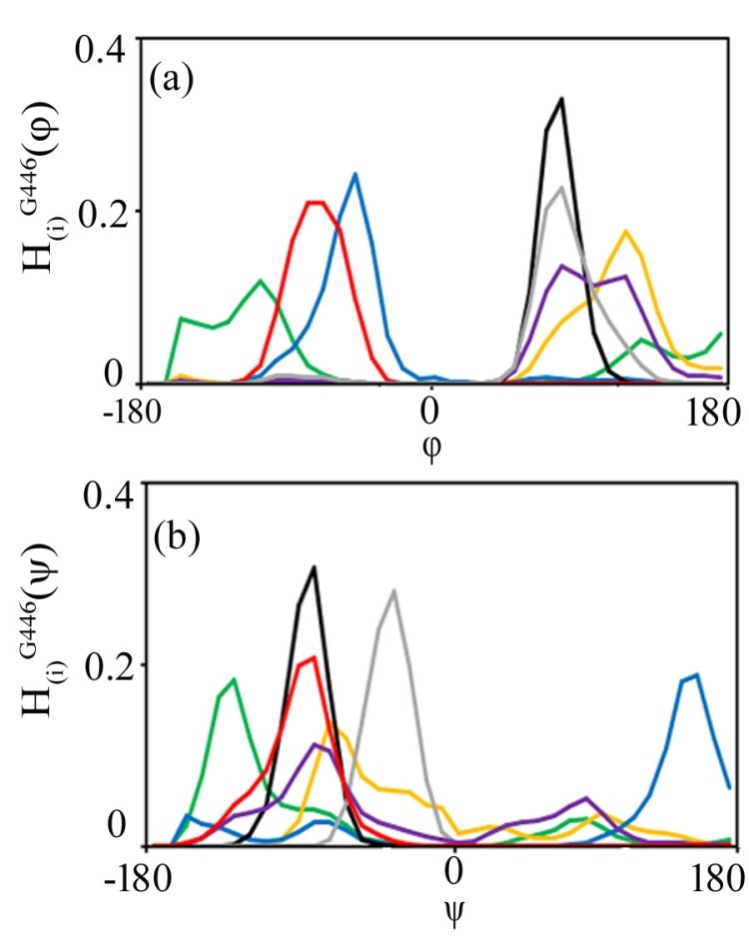

Fig. S2:

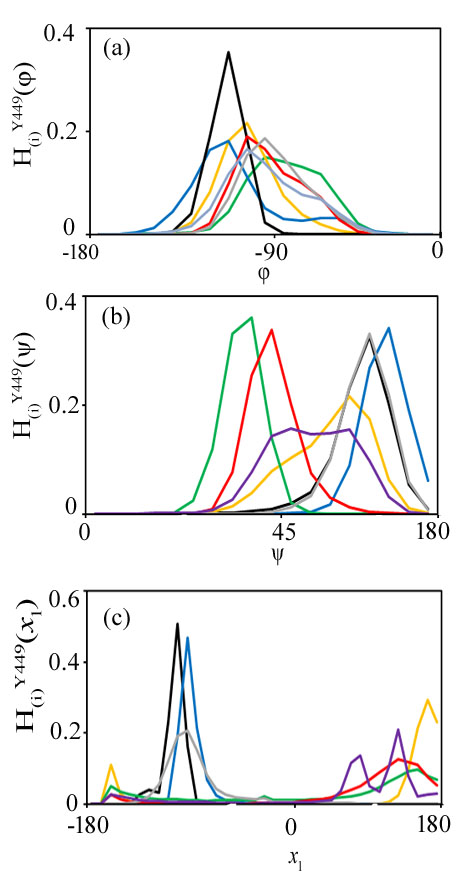

Fig. S3:

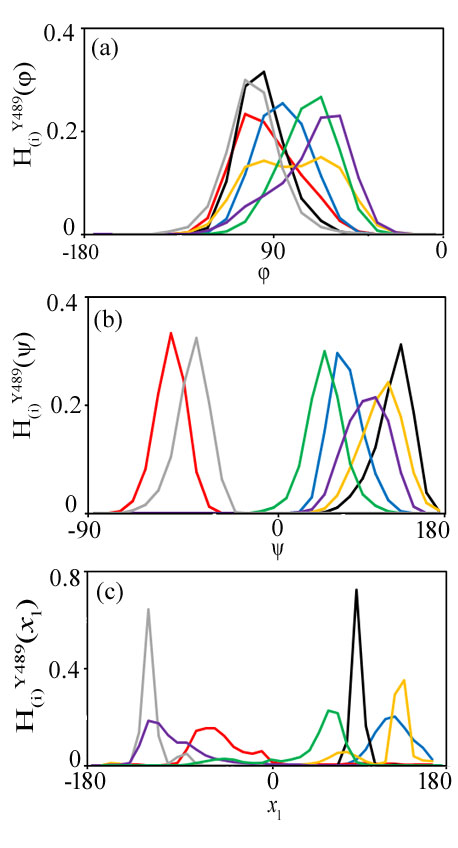

Fig. S4:

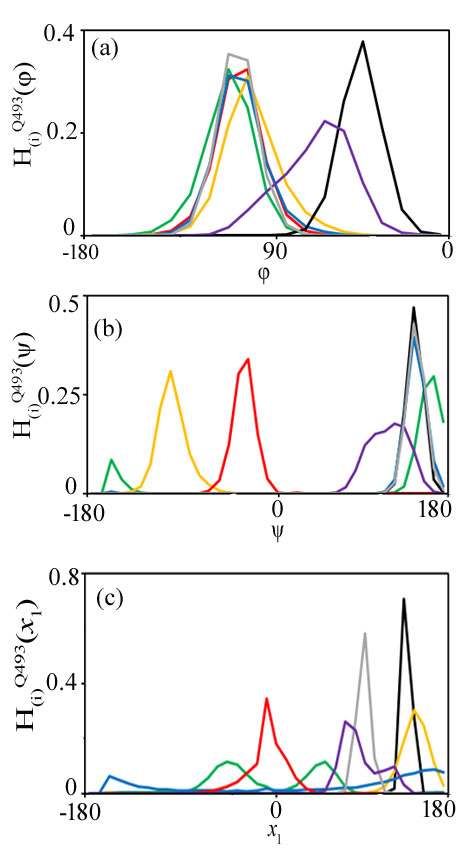

Fig. S5:

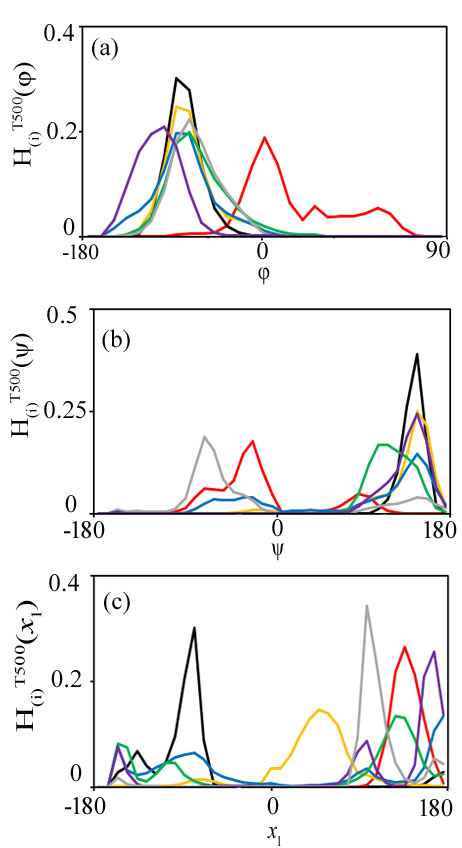

Fig S6

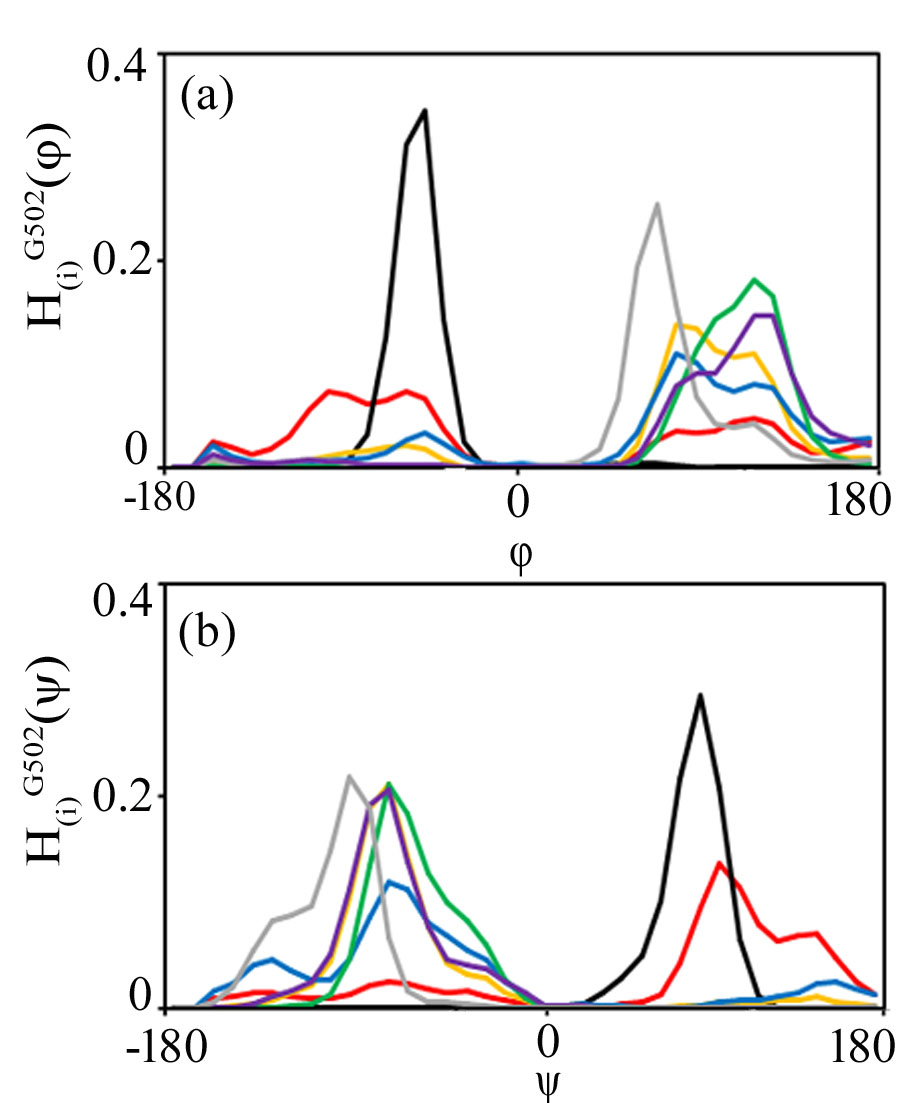

Fig. S7

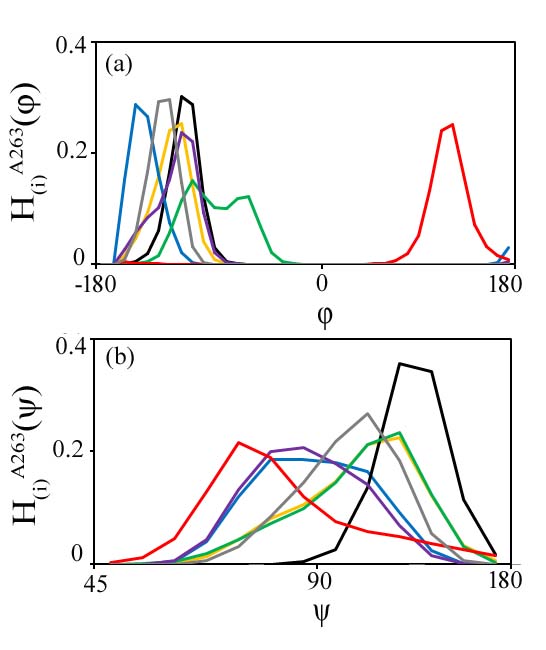

Fig S8

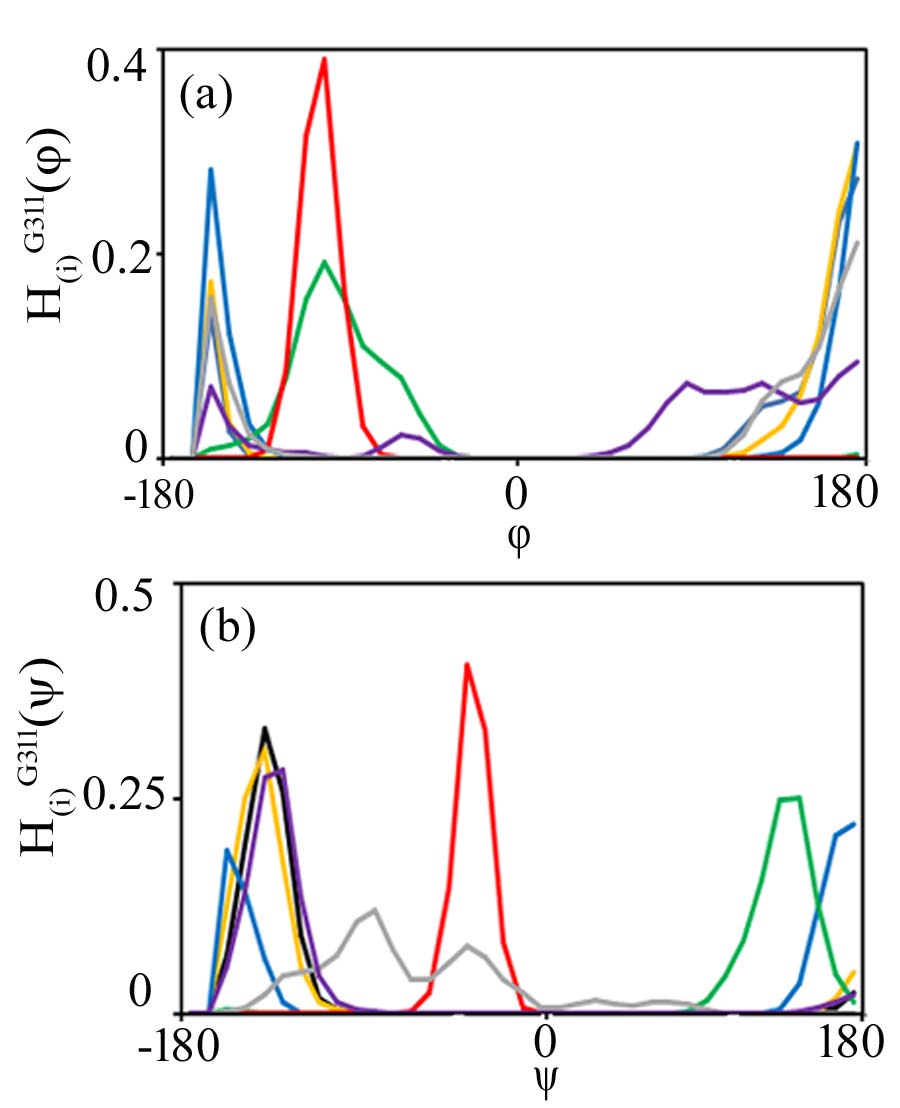

Fig S9:

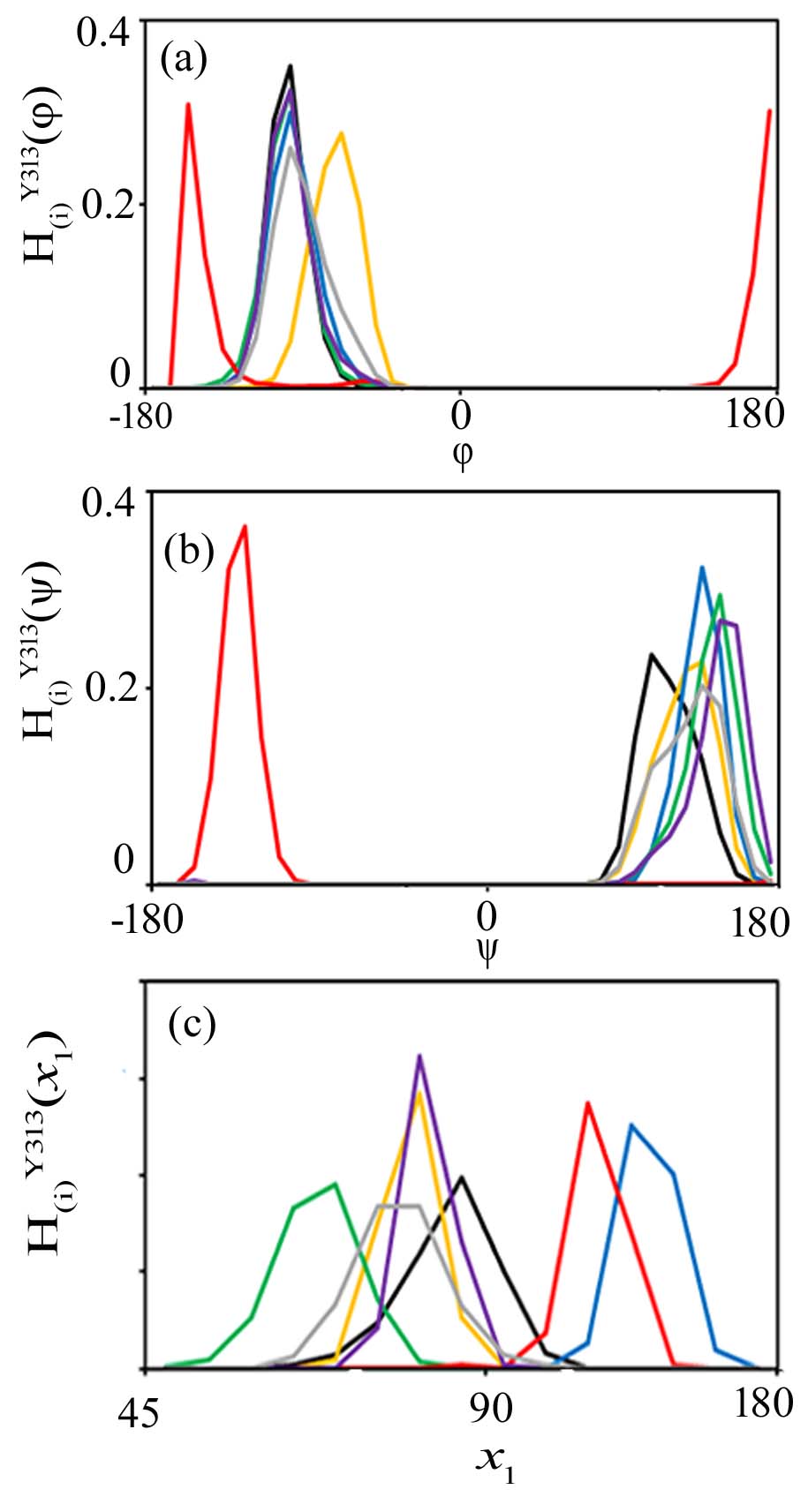

Fig. S10:

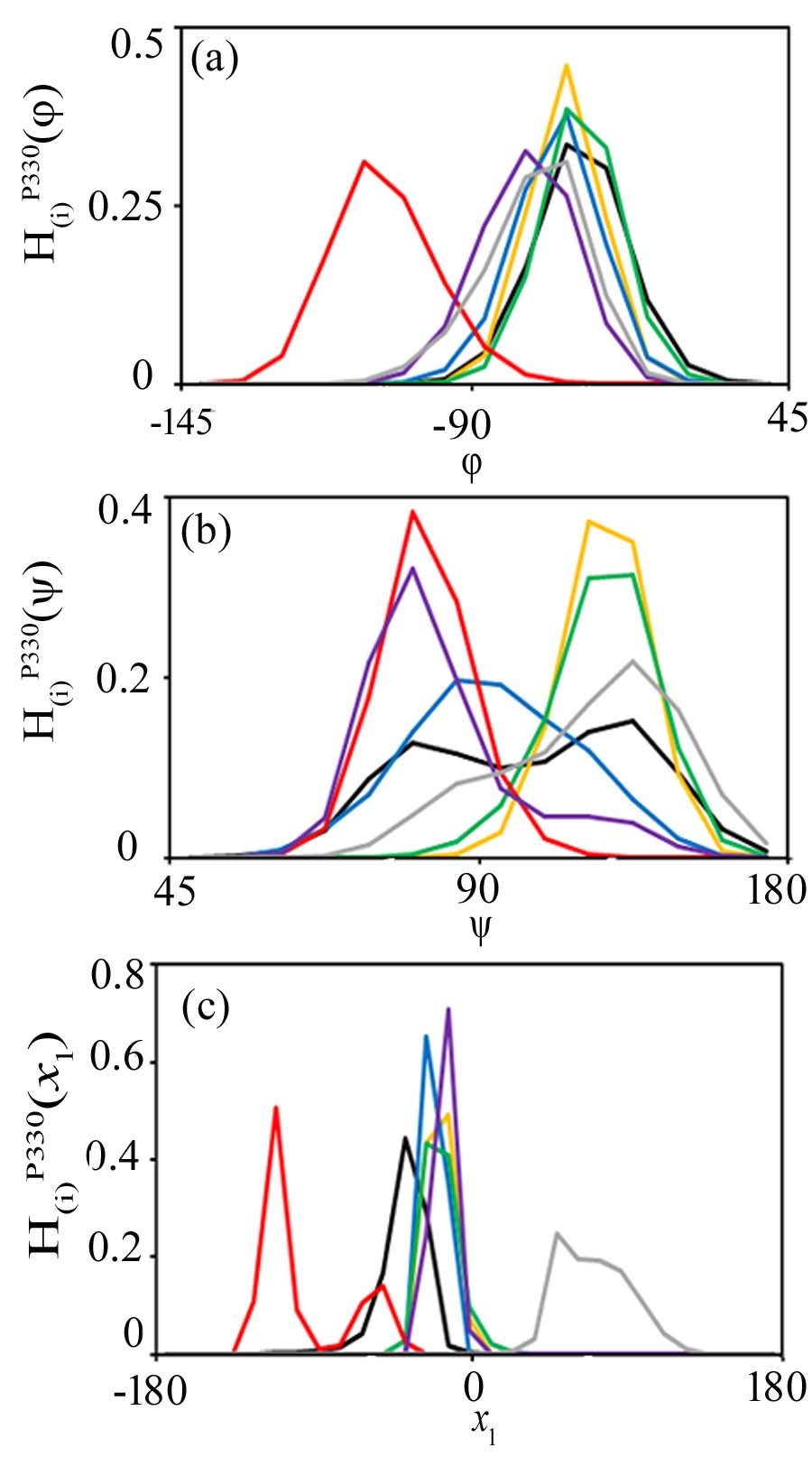

Fig S11:

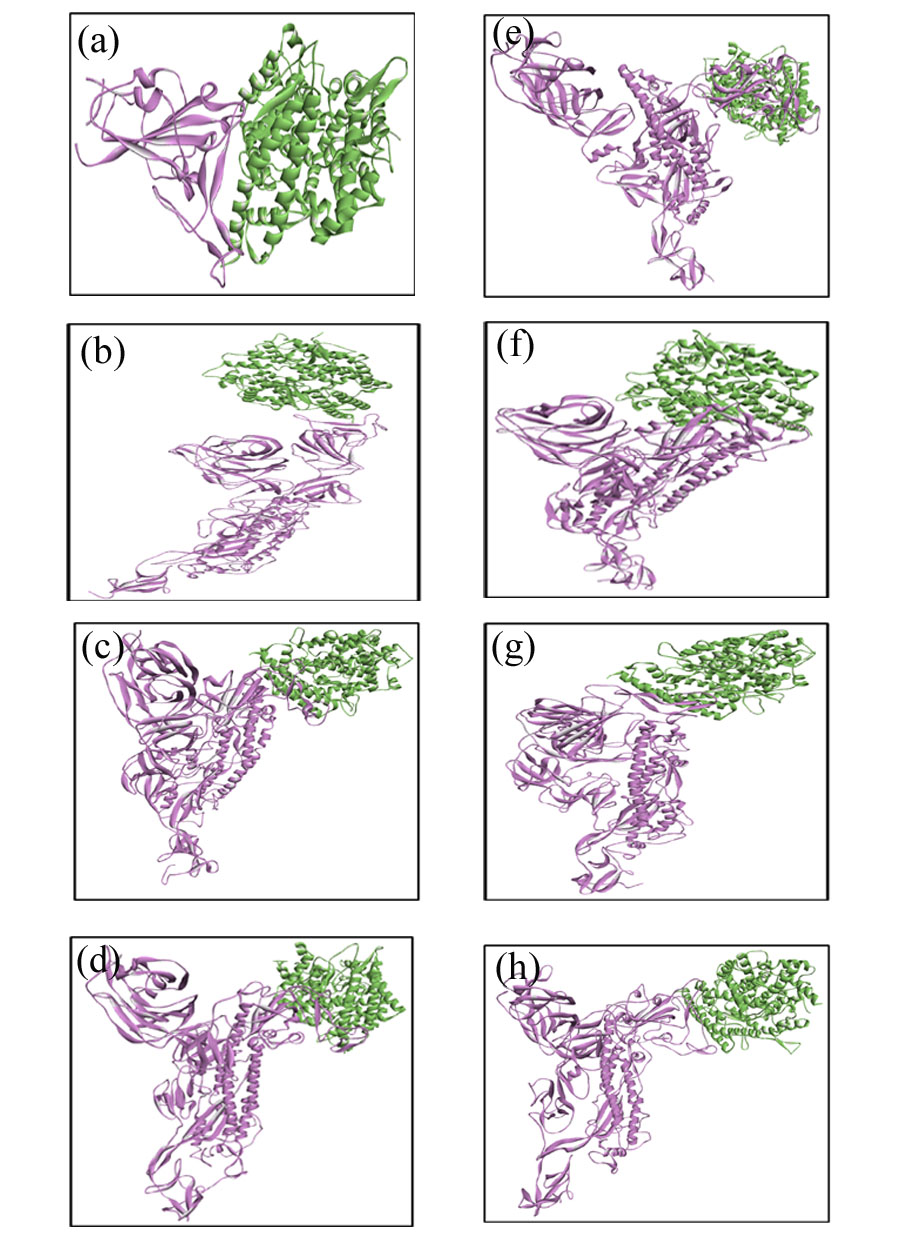
